# Reconstructing visual illusory experiences from human brain activity

**DOI:** 10.1101/2023.06.15.545037

**Authors:** Fan Cheng, Tomoyasu Horikawa, Kei Majima, Misato Tanaka, Mohamed Abdelhack, Shuntaro C. Aoki, Jin Hirano, Yukiyasu Kamitani

## Abstract

Visual illusions provide significant insights into the brain’s interpretation of the world given sensory inputs. However, the precise manner in which brain activity translates into illusory experiences remains largely unknown. Here we leverage a brain decoding technique combined with deep neural network (DNN) representations to reconstruct illusory percepts as images from brain activity. The reconstruction model was trained on natural images to establish a link between brain activity and perceptual features and then tested on two types of illusions: illusory lines and neon color spreading. Reconstructions revealed lines and colors consistent with illusory experiences, which varied across the source visual cortical areas. This framework offers a way to materialize subjective experiences, shedding new light on the brain’s internal representations of the world.

## Introduction

Visual illusions occur when perception of the world dissociates from sensory inputs. These illusions have been used to understand how the brain creates internal representations of the world. Physiological and neuroimaging studies have provided evidence of neural responses associated with illusory features. At the level of individual neurons in the visual cortex, some neurons exhibit similar responses to both actual and illusory attributes, suggesting the presence of a common neurobiological processing mechanism (Heydt et al., 1984; Peterhans and Heydt, 1989; Grosof et al., 1993; Lee and Nguyen, 2001; Ramsden et al., 2001; Roe et al., 2005; Sáry et al., 2007; Pan et al., 2012; Cox et al., 2013; Pak et al., 2019; Saeedi et al., 2022). On a broader scale, differential brain activity in some visual areas correlates with the perception of illusory attributes (Sasaki and Watanabe, 2004; Seghier and Vuilleumier, 2006; Cornelissen et al., 2006; Knebel and Murray, 2012; Kok and de Lange, 2014; Kok et al., 2016; Ho and Schwarzkopf, 2022), and brain activity patterns can be classified according to differences in illusory experiences (Hong and Tong, 2017; Gerardin et al., 2018). However, despite these insights, the precise impact of these neural responses on overall perceptual experience remains elusive. Elucidating how the population activity of visual cortical areas translates into the exact content of an illusory experience is essential to fill a critical gap in our understanding of how brain activity represents perceptual experience.

We address this issue by reconstructing illusory percepts as images from brain activity at different levels of processing in the visual cortex. Recent decoding and reconstruction techniques have utilized deep neural network (DNN) representations translated from brain activity to enable the reconstruction of arbitrary stimulus images (Miyawaki et al., 2008; Horikawa and Kamitani, 2017; Shen et al., 2019a,b). These techniques have also facilitated the reconstruction of subjective content, such as mental imagery and attention-modulated perception, by using the same model that was trained on stimulus perception (Shen et al., 2019a; Horikawa and Kamitani, 2022). Reconstruction provides a coherent representation of visual experience, encoded by neural population patterns and those mapped to DNN representations, which can be internally modulated or generated to counter sensory inputs. We hypothesize that an illusory stimulus would produce brain activity similar to that induced by a stimulus reflecting the subjective appearance of the illusion at specific stages of visual processing. This brain activity could be translated, or decoded, into DNN representations, which could then be converted into an image that exhibits the illusory attribute absent in the original stimulus.

## Methods

### Illusory stimuli and the reconstruction model

We tested this idea using representative line and color illusions — illusory lines induced by offset-gratings and neon color spreading (Fig. 1A; fig. S1; see Materials and Methods). Illusory lines were produced by shifted line gratings (Schumann, 1900; Soriano et al., 1996). We used a total of six configurations of the inducer (0*^◦^*, 90*^◦^*) and illusory orientations (0*^◦^*, 45*^◦^*, 90*^◦^*, and 135*^◦^*). Neon color spreading is an illusion where the color spreads out of the stimulus region, producing the percept of a transparent color surface (van Tuijl, 1975; Bressan et al., 1997). We used two versions of neon color spreading: the Ehrenstein configuration (Ehrenstein, 1941; Redies and Spillmann, 1981), where the color is restricted to the line regions but appears to spread out to form a circular surface, and the Varin configuration (Varin, 1971), where the color is restricted to the wedge regions but appears to spread out to form a rectangular surface. For both illusions, we prepared control images that weakened or abolished the illusory percepts, and positive control images that mimicked the illusory percepts with real lines or a uniform color surface.

**Figure 1.**
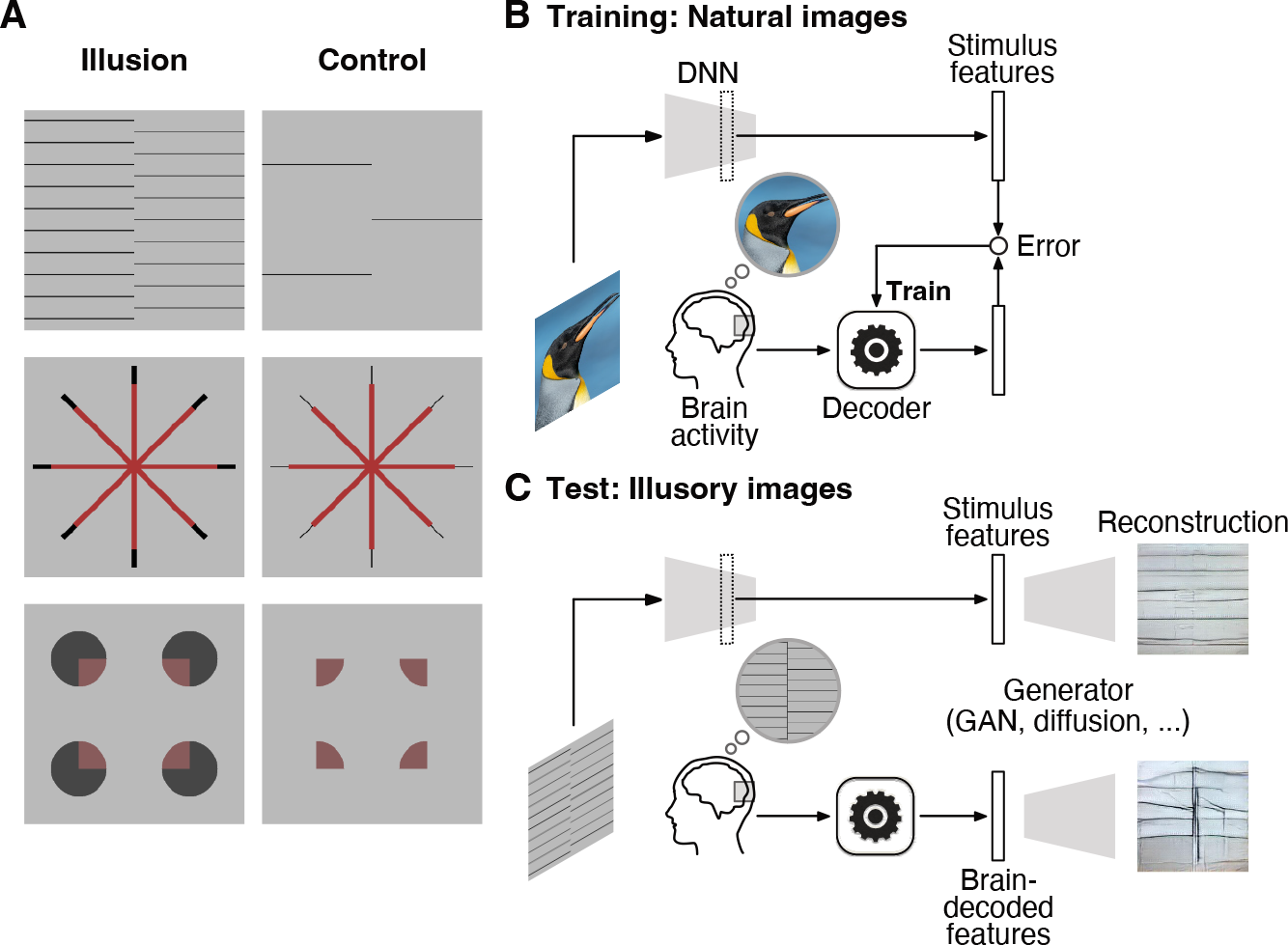
Illusory stimui and image reconsturction procedure. (**A**) Example images of the illusion (left) and control (right) conditions: an illusory line induced by offset-gratings (top), the Ehrenstein (middle) and Varin (bottom) configurations for neon color spreading. (**B**)Training. The stimulus features of natural images were extracted with a DNN pre-trained for object recognition. Decoders were trained to predict the stimulus (DNN) features from fMRI responses to the same images. (**C**) Testing. Illusory images were presented together with control and positive control stimuli. The stimulus features of a test stimulus or the DNN features decoded from fMRI responses to the test stimulus were passed to a pre-trained generator for reconstruction.

We adopted reconstruction models that consisted of DNN feature decoders and an image generator (Fig. 1, B and C). The DNN feature decoders were similar to those used in our previous studies (Horikawa and Kamitani, 2017; Shen et al., 2019a). We employed the unit activations of a feedforward convolutional neural network (Krizhevsky et al., 2012) as the target of decoding (Fig. 1B). The DNN feature decoders were trained on fMRI brain activity elicited by natural images of objects, material, and scenes including those added for this study (3,200 images in total; Materials and Methods). We used the fMRI signals of seven subjects from the visual cortex (VC), which covered both the early areas and the ventral object-responsive areas (see Materials and Methods). We assumed that most natural images would induce perceptions that closely mirror the physical features of the image (veridical perception) and that the trained decoders could adequately represent the mapping between brain activity and perceptual features, even without explicit information about subjective appearances.

At the test stage, we used the fMRI dataset measured for this study where the seven subjects were shown the illusory images as well as the control and positive control images in sequences interleaved with natural images (Fig. 1C; see Materials and Methods for exclusion criteria and missing data). Each test image was flashed at 0.625Hz for 8 seconds in each trial and it was repeated across twenty trials. Using the trained DNN feature decoders (Fig. 1B), we obtained the decoded features from each single-trial brain activity of the test dataset (Fig. 1C). The decoded features were then input to a generator, which had been trained on a large natural image dataset to convert the stimulus DNN features of an image back to the original image. The stimulus DNN features of the test illusory images were also given to the generator to see if the involvement of brain activity is critical for the reconstruction of illusory features. We primarily used a generator based on a generative adversarial network (GAN) (Dosovitskiy and Brox, 2016; VanRullen and Reddy, 2019), which we had decided on prior to data collection. However, we also tested additional generators based on diffusion methods (Ho et al., 2020; Dhariwal and Nichol, 2021; Yang et al., 2023), and pixel optimization (Shen et al., 2019a). While the three generators produced reconstructed images with different flavors, they yielded qualitatively similar results in terms of the visual features of interest. The chosen method was able to reconstruct natural images (fig. S2) with a quality that was on par with our previous study (Shen et al., 2019a).

## Results

### Reconstructed images

First, we confirmed that our reconstruction pipeline did not create spurious lines or colors congruent with the illusory percepts. We show reconstructions derived from stimulus DNN features for representative configurations (Fig. 2, “Stimulus features”). In all configurations, the reconstructions with stimulus features alone did not exhibit illusory components, even in the presence of noise (see fig. S3 for results from stimulus features plus the noise; see Materials and Methods). Although some DNN models can be trained to represent illusory appearances without the involvement of brain activity (Watanabe et al., 2018; Lotter et al., 2020; Gomez-Villa et al., 2020; Sun and Dekel, 2021), the DNN model we used as the target for feature decoding is a feedforward convolutional neural network trained for object classification (Krizhevsky et al., 2012), and thus is unlikely to represent contextual features like illusory line and color. In fact, we found that individual units of the DNN representation did not show orientation or color tuning shared between real and illusory features (figs. S4 and S5). Thus, our reconstruction model itself seems to translate visual information represented in brain activity following the coding rules for veridical perception.

**Figure 2.**
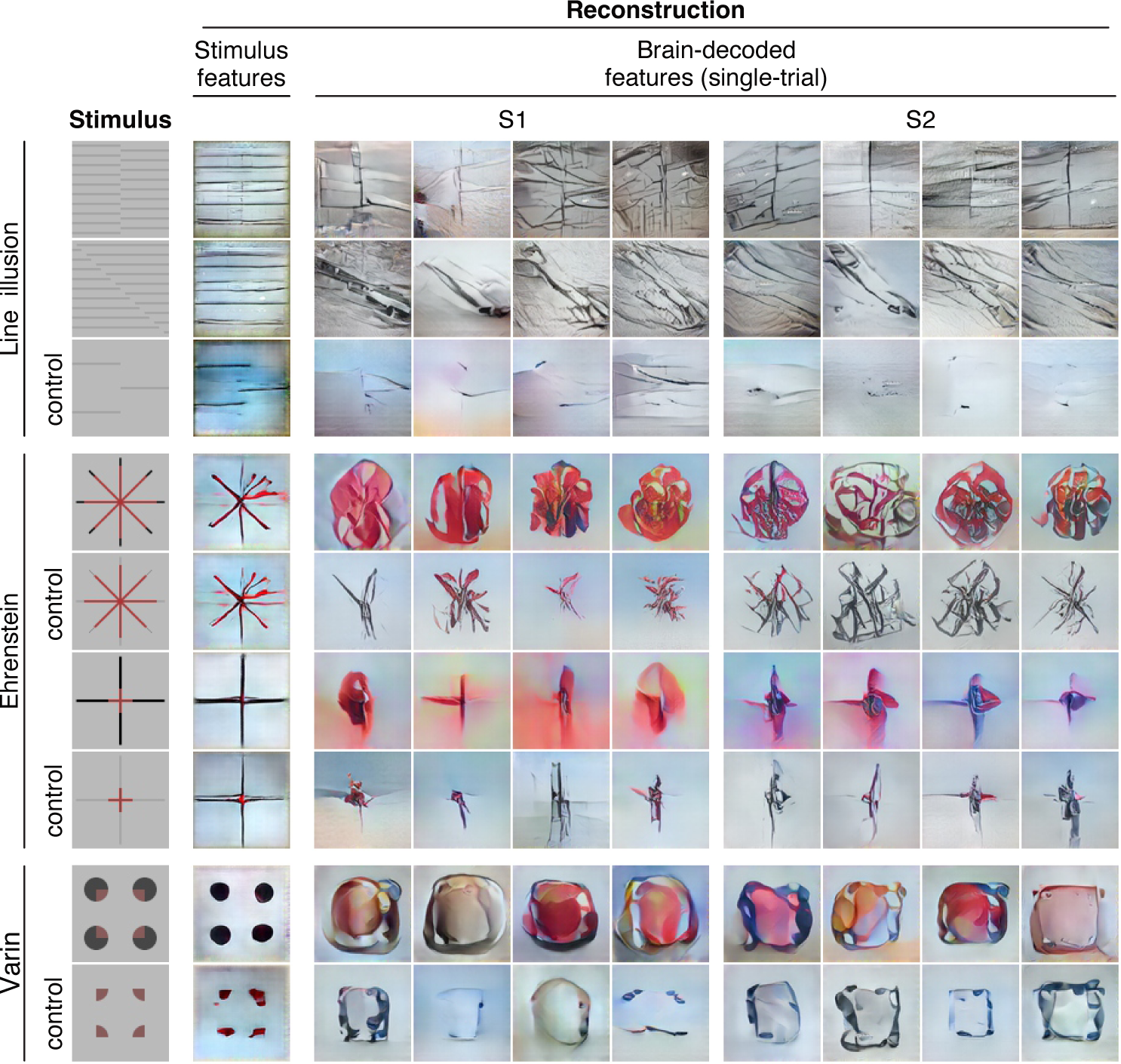
Reconstructions of illusory and control images. Reconstructions from stimulus features and from brain-decoded features are shown for two representative subjects (S1, S2). Reconstructions from brain-decoded features were produced from single-trial (8-sec) fMRI signals in the whole visual cortex (VC). Representative reconstructions from four different trials are shown for each subject.

Building on these findings, we then examined the reconstructions with DNN features decoded from single-trial brain activity in the whole visual cortex (VC) for each stimulus image (Fig. 2, right panel; two representative subjects and trials; see figs. S6–S9 for results of others and fig. S10 for results of other generators). In the line illusions, the reconstructions contained line components of the illusory and the inducer orientations. The illusory orientation often appeared more prominent than the inducer orientation, and this effect was not limited to the specific image region where the illusory line was actually perceived. In the control condition, where there were fewer grating lines to weaken the illusory percept, the line components of the illusory orientation were not as prominent.

In the Ehrenstein configuration of neon color spreading, the reconstructions exhibited a more extensive colored region compared to the control, where a line-width gap was introduced to eliminate the illusory percept. For the Varin configuration, the control image was designed to suppress the color spreading but not the contour or shape component. The reconstructions from the illusory and control conditions showed a contour-like intensity profile, but color spreading was more pronounced in the illusory configuration. In both the Ehrenstein and Varin configurations, the outer inducer parts were often poorly reconstructed, likely due to the selective flashing of color regions primarily located in more central positions. The lower resolution of the peripheral representation may also contribute to the inferior reconstructions observed in the periphery.

### Quantitative analyses of illusory lines across multiple brain areas

To quantify the reconstructed illusory lines (Fig. 3A), we detected the most apparent line orientation in each single-trial reconstruction using the Radon transform (Jafari-Khouzani and Soltanian-Zadeh, 2005). We calculated Radon projections for line areas traversing the center of an image at each orientation. A prominent line would cause a significant change in the projection value across parallel line areas at the line’s orientation. Thus, the orientation with the largest variance was defined as the principal orientation in each reconstruction (see Materials and Methods). The distribution of the principal orientations is shown in Figure 3B by pooling single-trial reconstructions from VC for stimulus images with a 90*^◦^*-difference between the illusory and inducer orientations (*n* = 275 trials that survived the exclusion criteria from seven subjects; see fig. S11A for results of each subject). This distribution had a bimodal peak at the illusory and inducer orientations, with 61.1% of principal orientations closer to the illusory than the inducer orientation. The configurations of a 45*^◦^*-difference showed a similarly high closer-to-illusory proportion (65.2%; fig. S11, A and B).

**Figure 3.**
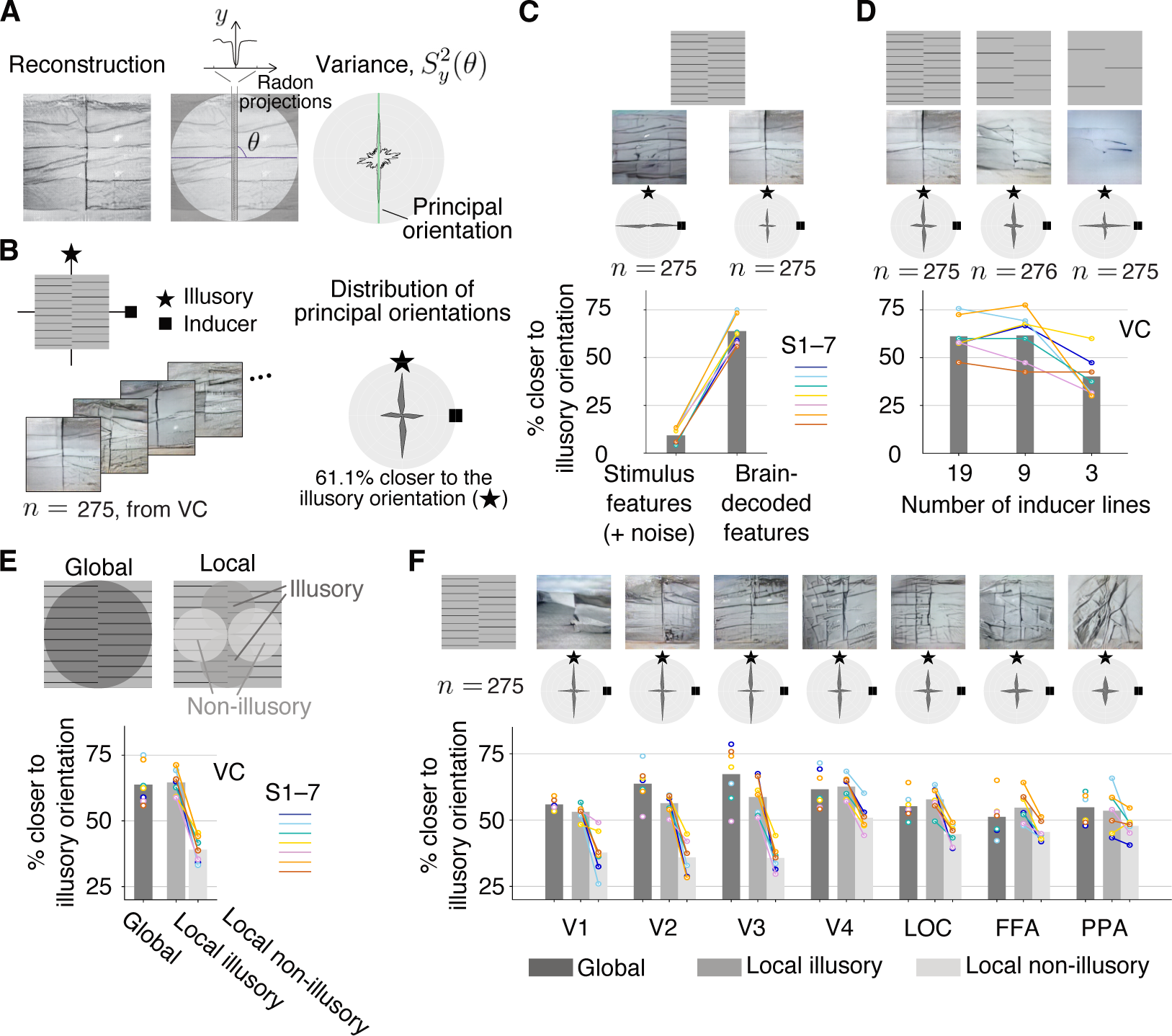
Evaluation of line illusion reconstructions. (**A**) Principal orientation detection. The orientation with the largest variance in Radon projections across line positions was identified as the principal orientation in an image. (**B**) Distribution of principal orientations in single-trial reconstructions from VC (results for seven subjects and all 90*^◦^*-difference configurations are pooled, totalling *n* samples; bin size = 15*^◦^*). An illusory orientation (star) and an inducer orientation (square) are shown for reference. (**C**) Comparison of reconstructions from stimulus features with added noise and brain-decoded features of VC. (**D**) Comparison of reconstructions with the different numbers of inducer lines. (**E**) Local presence of illusory orientation in reconstructions. (**F**) Comparison of reconstructions from individual visual areas. Reconstruction examples are from single-trial brain activity in VC (except in F) of subject S2. The polar plots show the distributions of the principal orientations pooled across all subjects and all 90*^◦^*-difference configurations. The bar graphs indicate the proportions of principal orientations closer to the illusory than to the inducer orientation, pooled for all subjects and configurations. Color circles and lines indicate individual subjects.

Using these methods, we compared the reconstructions from brain-decoded features of VC with those from stimulus features (noise added to match the decoded features from each subject; see Materials and Methods). Consistent with representative reconstructions (Fig. 3C, top panel; see other reconstructions in figs. S3 and S6), the principal orientations with stimulus features were distributed mostly around the inducer orientation, unlike the bimodal distribution found with brain-decoded features (Fig. 3C, middle panel; all 90*^◦^*-difference con-figurations and subjects pooled). The closer-to-illusory proportion for stimulus features (plus noise) was near zero (Fig. 3C, bottom panel; all configurations pooled in each subject; one-sided z-test, *p* < 0.01 in 7/7 subjects). These results further confirm that the reconstructed illusory lines were derived from brain activity, not from the analysis pipeline itself.

Additionally, we investigated the effect of the number of inducer lines. Prior research has shown that decreasing the number of inducer lines weakens illusory percepts (Soriano et al., 1996). The reconstructions (Fig. 3D, top panel; additional examples in fig. S7) and distributions of the principal orientations (Fig. 3D, middle panel; all 90*^◦^*-difference configurations and subjects pooled; see results of each subject in fig. S12) indicate a grad-ual reduction in the strength of the illusory lines as the number of inducer lines is decreased. The closer-to-illusory proportion was similar at line numbers 19 and 9 and then decreased at line number 3 (Fig. 3D, bottom panel; all 90*^◦^*-difference configurations and subjects pooled). Thus, the reconstructions appear to reflect the illusory appearance manipulated by the number of inducer lines.

The illusory line is perceived at the abutting portion of the inducer gratings. However, the method used above for detecting the principal orientation fails to capture this locality of the illusory percept. To examine the local presence of the illusory orientation in reconstructions, we analyzed local image regions and detected the principal orientation separately at 1) illusory regions where the illusory line was expected to be seen and at 2) non-illusory regions where only inducer lines were expected to be seen (Fig. 3E, top panel). We calculated the closer-to-illusory proportions at each local region as well as the global region from the previous analysis (Fig. 3E, bottom panel; all configurations pooled in each subject). While the closer-to-illusory proportions for the local illusory regions were similar to those for the global region, the proportions for local non-illusory regions were significantly lower (one-sided z-test for proportion, *p* < 0.01 in 7/7 subjects; see fig. S11C for results of 90*^◦^*- and 45*^◦^*-difference configurations).

Reconstructions can be obtained using fMRI activity in individual visual areas of the visual cortex (V1–V4, lateral occipital complex [LOC], fusiform face area [FFA], parahippocampal place area [PPA]; see Materials and Methods). In Figure 3F, we show examples of the reconstructions from individual areas (top panel; see fig. S13 for others), the distributions of principal orientations (middle panel; pooled across 90*^◦^*-difference configurations and subjects; see fig. S14 for results of 90*^◦^*- and 45*^◦^*-differences from each subject), and the closer-to-illusory proportions for the global, local illusory, and local non-illusory regions (bottom panel; all configurations pooled in each subject). Overall, V1–V3 tended to show faithful reconstructions of illusory and inducer lines. V4 and the higher visual areas showed less localized illusory lines and poorly reconstructed inducer lines. The strength of the illusory orientation component for the global regions peaked around V2–V4 (Fig. 3F, bottom panel). The difference between local illusory and non-illusory regions, which indicates the consistency with the local illusory percept, was large around V1–V3 (one-sided z-test for proportion, *p* < 0.05 in 5/7 subjects at V1, 7/7 at V2, V3, and V4, 6/7 at LOC, 4/7 at FFA, 3/7 at PPA). The overall global and local trends in reconstructions of illusory images were found to mirror the trends seen for the positive control images, though the actual strength of the illusory line was weaker than that of the real line in V1–V3 (fig. S15). The strength of illusory orientation components (indicated by global closer-to-illusory proportions) in V1–V4 and LOC seem to reflect the strength of subjective line perception manipulated by the number of lines (fig. S16). The results suggest that the local representations of illusory lines are formed in early visual areas and that illusory and real lines are similarly represented across visual areas.

### Quantitative analyses of illusory color across multiple brain areas

We next produced reconstructions for neon color spreading from individual visual areas as well as VC (Fig. 4, A and B; see figs. S8, S9, S17, and S18 for additional examples). Reconstructions from the illusion condition of the Ehrenstein configuration exhibited red regions across areas, similar to the positive control condition, with lower areas more accurately depicting the sizes of red regions and inducer lines (Fig. 4A). However, in the control condition, where a gap in line width was introduced to abolish or weaken illusory color spreading, the color was largely absent in the reconstructions, even though the illusion and control stimuli included the same red regions. Reconstructions from the Varin illusion condition exhibited similarity to the positive control condition only in middle-to-higher areas, showing broad red regions (Fig. 4B). The color was barely present at V1–V3, while faithful reconstructions of the real color could be produced from V2 and V3 in the positive control condition. The inducer regions were poorly reconstructed, even in lower areas, in the illusion and the positive control conditions. Although square-like outlines were seen in the reconstructions from the lower areas, they did not coincide with the illusory square shape but seemed to enclose the entire stimulus region. In the control condition, the color was almost absent: square outlines were reconstructed more clearly from the lower areas and appeared to align the illusory square without color spreading.

**Figure 4.**
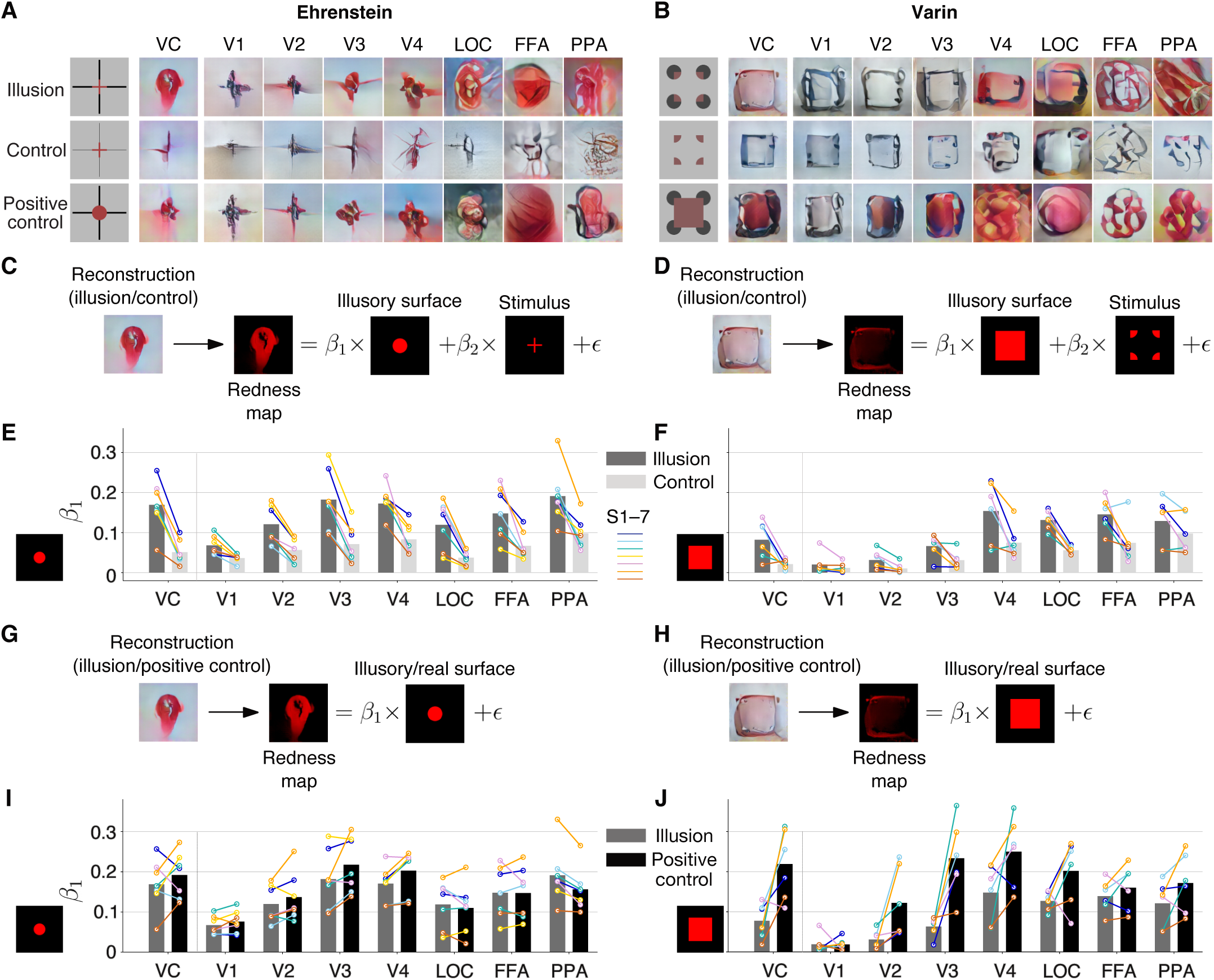
Evaluation of neon color spreading reconstructions. (**A** and **B**) Representative single-trial reconstructions of the illusion (top), control (middle), and positive control (bottom) conditions for Ehrenstein from subject S1 (A) and for Varin from subject S2 (B). (**C** and **D**) Illustration of regression analysis for comparing the illusion and control conditions for Ehrenstein (C) and Varin (D). The redness map of a reconstructed image was fitted by those of the illusory surface (expected region of color filling-in) and the stimulus. (**E** and **F**) Comparison of the illusory surface coefficient values between illusion and control conditions for Ehrenstein (E) and Varin (F). Results for all configurations (sizes and numbers of lines) and seven subjects are pooled for Ehrenstein. Results for six subjects are pooled for Varin. Color lines indicate the results of individual subjects. (**G** and **H**) Illustration of regression analysis for comparing the illusion and the positive control conditions for Ehrenstein (G) and Varin (H). The redness map of a reconstructed image was fitted by that of the illusory or real surface. (**I** and **J**) Comparison of the illusory surface coefficient values between the illusion and the positive control conditions for Ehrenstein (I) and Varin (J), pooled as in (E) and (F).

To quantify color spreading, we performed regression analysis on the pixel color values in each reconstructed image (Fig. 4, C and D; see Materials and Methods). We created the redness (saturation) maps from the original RGB values of reconstructed and stimulus images. Additionally, we prepared the redness maps for the expected illusory surface regions. The profile of the redness map from a reconstruction was fitted by those of the expected illusory surface region and the stimulus (regressors with coefficients 1 and 2, respectively). The illusory surface regressor was shared between the illusion and the corresponding control conditions for comparison. The illusory surface coefficient (1) was used as the measure for illusory color reconstruction. Regression coefficients were calculated for all individual trials (reconstructions) and pooled across different configurations (sizes and numbers of lines for Ehrenstein) in each subject and brain area.

For Ehrenstein, the illusory surface coefficient was generally higher in the illusion condition than in the control condition (Fig. 4E; one-sided t-tests in individual sub-jects, *p* < 0.05 in 7/7 subjects at VC, 5/7 at V1, 7/7 at V2, V3, V4, LOC, and FFA, and 6/7 at PPA; see fig. S19A for results with different stimulus configurations). V2–V4 and higher areas appear to show robust illusion effects (see fig. S20 for results with individual trials in each subject). For Varin, the illusory surface coefficient was greater in the illusion than in the control condition in mid-to-higher areas but the effects in individual subjects were less robust than those with Ehrenstein (Fig. 4F; one-sided t-tests in individual subjects, *p* < 0.05 in 3/6 subjects at VC, 2/6 at V1, V2, V3, and 3/6 at V4, 4/6 at LOC, 3/6 at FFA, 1/6 at PPA; see fig. S21 for results with individual trials in each subject).

Similarly, we performed regression analysis in which the profile of the redness map from a reconstruction was fitted by the real or illusory color surface for the illusion and the positive control conditions (Fig. 4, G and H). Whereas surface coefficients were comparable between the illusion and the positive control conditions across brain areas for Ehrenstein, lower surface coefficients were observed in the illusion compared to the positive control condition in low-to-middle areas for Varin (Fig. 4, I and J; see figs. S22 and S23 for results with individual trials in each subject). Additionally, large-sized Ehrenstein configurations tended to show lower illusory surface coefficients for the illusion than the positive control condition in the low-to-mid brain areas (fig. S19B). Thus, the strength of illusory color reconstruction across brain areas may depend on the spatial extent of filling-in as well as the stimulus configurations.

## Discussion

We have demonstrated the reconstruction of illusory percepts as images from single-trial brain activity, using the computational model that learns the coding scheme linking non-illusory stimulus-induced perception and brain activity. The reconstructions obtained from the visual cortex resembled the illusory percepts, to the point where the relevant attributes were perceptually recognizable. It is important to note that neither the stimulus features of the illusion-inducing stimuli nor the brain-decoded features of the control images produced such reconstructions. One notable advantage of our approach is the ability to externalize mental contents, going beyond the conventional method of testing qualitative hypotheses. By representing mental contents in a manner comprehensible to others, we offer a means of sharing and understanding subjective experiences. However, the current methods have limitations including the lack of spatial resolution, particularly for the peripheral vision, resulting in a poor reconstruction of the inducers. Additionally, the reconstructed images are distorted presumably due to inherent biases in the reconstruction model.

The reconstructions from individual areas unveiled the strength of illusory representation and the extent to which it is shared with real stimuli at different processing stages. Specifically, regarding illusory lines, our results align with previous findings (Heydt et al., 1984; Peterhans and Heydt, 1989; Grosof et al., 1993; Lee and Nguyen, 2001; Ramsden et al., 2001; Seghier and Vuilleumier, 2006; Sáry et al., 2007; Knebel and Murray, 2012; Pan et al., 2012; Cox et al., 2013; Pak et al., 2019), showing the involvement of low-level areas and LOC in processing both illusory and real lines. Our reconstructions further revealed that local illusory lines, which accurately reflected perceptual experiences, were better represented in areas V1–V3, while central illusory lines tended to dominate in higher areas. With regard to illusory color, previous research has produced mixed findings concerning its represen-tation in the low and middle areas (Sasaki and Watanabe, 2004; Cornelissen et al., 2006; Hong and Tong, 2017; Gerardin et al., 2018). Our results, however, show the participation of multiple areas in this process, extending even to higher areas within the ventral cortex. Specifically, for the Ehrenstein configuration, we observed that illusory color representations spanned from low to high areas, with overlaps with real color representations. In contrast, in the Varin configuration, we found illusory color representations primarily in mid-to-high areas, whereas early areas manifested a more pronounced representation of real color as compared to illusory color.

Our findings suggest that neural representations of illusory percepts vary not only between different visual attributes such as line and color, but also among configurations for the same attributes. It is increasingly clear that no single brain region can entirely account for subjective experience; rather, multiple neural mechanisms across various areas are likely implicated in the processing and interpretation of illusory stimuli. This nuanced understanding provides valuable insight into the neural representation of conscious experience. However, it is important to acknowledge that our methods primarily illuminate representations rather than processing or dynamics. To further our understanding, highresolution and multimodal brain measurements would serve as beneficial supplementary tools.

In conclusion, our reconstruction approach effectively bridges the gap between internal representations and their external manifestations. This method paves the way for novel explorations of, and communications about, the internal representations of the perceptual world that reside within the brain.

## Acknowledgments

We thank our laboratory team, especially Eizaburo Doi, Ken Shirakawa, and Yoshihiro Nagano, for their invaluable comments and suggestions on the manuscript. We want to extend a special thanks to our engineer, Mitsuaki Tsukamoto, who maintained our computing environment throughout this project. We gratefully acknowledge the assistance of the staff at the Kyoto University Institute for the Future of Human Society who ensured the smooth running of our experiments. Special thanks to all participants in our fMRI experiments without whom this research would not have been possible.

## Funding

Japan Society for the Promotion of Science (KAKENHI grant JP20H05705 and JP20H05954 to Y.K.); Japan Science and Technology Agency (CREST grant JPMJCR18A5 and JPMJCR22P3 to Y.K., and SPRING grant JPMJSP2110 to F.C.); New Energy and In-dustrial Technology Development Organization (Project JPNP20006 to Y.K.).

## Author contributions

Conceptualization: F.C., Y.K.; Data curation: F.C., M.T., S.C.A.; Formal analysis: F.C.; Funding acquisition: F.C., Y.K.; Investigation: F.C., T.H., M.T., M.A.; Methodology: F.C., T.H., K.M., M.T., H.J., Y.K.; Software: F.C., S.C.A.; Supervision: Y.K.; Visualization: F.C., Y.K.; Writing – original draft: F.C., Y.K.; Writing – review & editing: F.C., T.H., K.M., M.T., M.A., Y.K.

## Competing interests

Authors declare that they have no competing interests.

## Data and materials availability

The raw and preprocessed experimental data will be made public in OpenNeuro and figshare, respectively. The codes will be deposited in our repository (https://github.com/KamitaniLab).

## Materials and Methods

### Subjects

We collected fMRI data from seven healthy subjects, four males and three females (aged 25–36 years). All subjects provided informed consent prior to the experiment, with the study protocol having been reviewed and approved by the Ethics Committee of the Graduate School of Informatics at Kyoto University (approval no.: KUIS-EAR-2017-002). The subjects had normal or corrected-to-normal vision. Four subjects (S1–4) were the same as those in previous studies (Shen et al., 2019a; Horikawa and Kamitani, 2022). Therefore, we utilized their published data (data from the training session; available from https://openneuro.org/datasets/ds003430/versions/1.1.1 for subjects S1–3 and https://openneuro.org/datasets/ds001506/versions/1.3.1 for subject S4) as a subset of training data and collected additional training data. For subjects S5–7, training datasets were newly acquired with fewer repetitions in order to conserve resources (“fMRI Experiments: Training session”). For subjects S5 and S7, the same image set as subjects S1–4 was used, while for subject S6, 32 training images were replaced due to unpleasant object categories for the subject. Test datasets were newly acquired for all subjects though subject S4 lacked a few sessions of the test dataset (“fMRI Experiments: Test session”).

### Visual stimuli

#### Natural images

A total of 3,200 natural images were downloaded from the online image databases. We obtained 1200 object images from ImageNet (Russakovsky et al., 2015), 1000 material images from Flickr Material Database FMD (Sharan et al., 2014), and 1000 object or scene images from COCO (Lin et al., 2014). The images were cropped to a square at the center and resized to 500 × 500 pixels.

#### Illusory line images

We created six images with illusory lines induced by offset gratings (fig. S1A), and confirmed, prior to the fMRI experiments, that they induced clear line illusion for all subjects. Two gratings were used to fill the gray background image with a predetermined number of equally spaced black lines and a phase shift of half a cycle between the gratings. Hence, there were three parameters: the orientation of the illusory line (illusory orientation), the orientation of the inducer lines (inducer orientation), and the number of inducer lines. We set the number of inducer lines at 19, while varying the illusory orientations (0*^◦^*, 45*^◦^*, 90*^◦^*, and 135*^◦^*) and inducer orientations (0*^◦^* and 90*^◦^*). This resulted in six illusory line images, as the illusory and inducer orientations cannot be the same in a single image. We reduced the number of inducer lines to 9 and 3 for configurations with vertical or horizontal illusory orientation, respectively, creating four control images that induce weaker illusion perception. These parameters were based on previous behavioral studies on human subjects (Soriano et al., 1996), and adapted for our experiments. Additionally, we included ten corresponding positive control images with a real line drawn at the location where the illusory line was perceived.

#### Neon color spreading images

There are two types of neon color spreading images: the Ehrenstein configuration and the Varin configuration. For Ehrenstein (fig. S1B), we prepared four different versions with two sets of lines (four and two) and two sizes (small and large) of the colored portion. The luminance of the colored lines was set between the luminance values of the surrounding black lines and the background (gray) to meet the key requirement for transparency perception (Metelli, 1985; Nakayama et al., 1990). The colored lines and the surrounding black lines were connected in the same width, inducing color filling-in perception (transparent color disk). We constructed control images by reducing the width of the black lines, which created disconnected patterns and disrupted the color filling-in perception (Redies and Spillmann, 1981). In addition, we added positive control images with uniform color in the expected filling-in areas, while keeping the same black lines as in the illusion images.

For Varin (fig. S1C), the illusion image was composed of four disks, the centers of which could be connected to form a rectangle. Each disk was pieced by a colored 90◦ sector and a black Pacman. Similar to Ehrenstein, the luminance of the color was made between the luminance values of Pacman (black) and the background (gray). For control, we removed the black Pacman and retained only colored sectors to reduce the color filling-in effect (the main control image used in the analysis of Figure 4; left in the “Control” panel of fig. S1C), which shared the colored lines with the illusion image. Two other control images were created using only black Pacmans or disks. We also prepared two positive control images with uniform color in the rectangular region of interest. The one with the black packmen as in the illusion image was used for the analysis of Figure 4.

In total, we produced 12 images for Ehrenstein and 6 images for Varin. We selected red for the colored components due to its superior reconstruction quality compared to other colors (Shen, et al., 2019a). The saturation of the red color in each pattern was adjusted to ensure that all subjects clearly perceived color filling-in (0.8 for four-line and 0.7 for two-line frames of Ehrenstein, and 0.3 for Varin, respectively).

### fMRI Experiments

We performed image presentation experiments for two types of sessions: training and testing. All stimuli were rear-projected onto a screen in an MRI scanner bore with a luminance-calibrated liquid-crystal display projector. The stimulus images were displayed at the screen center with a size of 12*^◦^ ×* 12*^◦^* of visual angle on a gray background. We asked subjects to fixate on the center of the images cued by a circle of 0.3*^◦^ ×* 0.3*^◦^* of visual angle. Each subject used a custom-molded bite bar and/or personalized headcase from CaseForge, Inc. to reduce head motion during fMRI data collection. Multiple scanning sessions were performed to collect data for each subject. The total time span of data collection varied among the subjects: approximately 2 years for subjects S1–2, 4 years for subjects S3–4, and less than 1 year for subjects S5–7. Each consecutive session took a maximum of 2 hours, with each run taking 6-8 minutes. The subjects were free to rest adequately between runs or to terminate the experiment at any time.

#### Training session

A single set of training sessions consisted of presenting each of the 3,200 natural images once, resulting in 64 runs. Images from different training image sets (ImageNet, FMD, and COCO) were presented in separate runs. There were 32- and 6-sec rest periods at the beginning and end of each run, respectively. Each run contained 55 trials, with 50 trials of different images and five randomly interspersed repetition trials that showed the same image as the previous trial. Each image was flashed at 1 Hz during an 8-sec trial. There was no rest period between trials. To indicate the onset of a trial, the color of the fixation spot was changed to red 0.5 s before the trial and then switched back to white when the trial began. Subjects were instructed to perform a one-back repetition detection task, in which they were required to press the button immediately after a repeated image appeared. We asked the subjects to maintain steady fixation throughout the run and evaluated their alertness level using one-back task performance. We repeated one set of training sessions five times for subjects S1–4 (3,200 *×* 5 = 16,000 training samples) and two times for subjects S5–7 (3,200 *×* 2 = 6,400 training samples). Before analyzing the illusion test data, we confirmed that training data with fewer repetitions could produce a comparable model performance on an independent test dataset with natural image presentation (Shen et al., 2019a). The order of image presentation was randomly assigned across runs.

#### Test session

In the test session, we presented each of the 38 test images (10 illusory line images, 10 real line images, 12 Ehrenstein images, and 6 Varin images) 20 times. For subject S4, we only collected data for 32 test images because of the subject’s relocation before we decided to add the Varin configurations (6 images). We included the same number of randomly selected natural images to make the fMRI signal baselines comparable to the training sessions. The test session consisted of 40 runs. Similar to the training session, there were 32- and 6-sec rest periods at the beginning and end of each run, respectively. Each run contained 42 trials with 38 trials of different images (19 test images and 19 natural images) and four randomly interspersed repetition trials that showed the same image as the previous trial (subject S4 underwent fewer trials due to the fewer illusion images). To eliminate the after-effect from the previous test image, we inserted a natural image trial between every two illusion image trials. Otherwise, the presentation order was random in each run. Each image was flashed at 0.625 Hz during an 8-sec trial. For the neon-color illusion stimuli (Ehrenstein and Varin), only the colored portions were flashed on black lines or disks to enhance illusory percepts. There was no rest period between trials. The subjects performed a one-back repetition detection task based on the same cue from the fixation spot as in the training session.

### MRI acquisition

We collected fMRI data using a 3.0-Tesla Siemens MAGNETOM Verio scanner at the Kyoto University Institute for the Future of Human Society (formerly, Kokoro Research Center). An interleaved T2*-weighted gradient-echo echo-planar imaging (EPI) scan was performed to acquire functional images covering the entire brain (TR, 2000 ms; TE, 43 ms; flip angle, 80*^◦^*; FOV, 192 *×* 192 mm^2^; voxel size, 2 *×* 2 *×* 2 mm^3^ ; slice gap, 0 mm; number of slices, 76; multiband factor, 4). T1-weighted (T1w) magnetization-prepared rapid acquisition gradient-echo (MP-RAGE) fine-structural images of the entire head were also acquired (TR, 2250 ms; TE, 3.06 ms; TI, 900 ms; flip angle, 9*^◦^*; FOV, 256*×*256 mm^2^; voxel size, 1.0 *×* 1.0 *×* 1.0 mm^3^).

### fMRI data preprocessing

We performed the MRI data preprocessing through the pipeline of FMRIPREP (version 1.2.1). For the functional data of each run, we first estimated a BOLD reference image using a custom methodology of FMRIPREP. Then, data were motion-corrected using MCFLIRT from FSL (version 5.0.9) and slice time corrected using 3dTshift from AFNI (version 16.2.07), based on this BOLD reference image. Next, we co-registrated the corresponding T1w image using boundary-based registration implemented by bbregister from FreeSurfer (version 6.0.1). The coregistered BOLD time-series were then resampled onto their original space (2 *×* 2 *×* 2 mm^3^ voxels) with antsApplyTransforms from ANTs (version 2.1.0) using Lanczos interpolation. After obtaining the resampled BOLD time series, we first shifted the time series by 4 seconds (2 volumes) to compensate for hemodynamic delays, and then regressed out nuisance parameters from each voxel’s time series of each run, including a constant baseline, a linear trend, and temporal components proportional to the six motion parameters calculated during the motion correction procedure (three rotations and three translations). We created single-trial data samples by reducing extreme values (beyond *±*3 SD for each run) of the time series and averaging within each 8-sec trial (four volumes).

### Data exclusion criteria

We fixed the data exclusion criteria before fMRI data collection and finished exclusion before proceeding to the main analyses. First, runs with low performance (hit rate ≤ 50 %) in the one-back repetition detection task were excluded. This step was to discard the scanned data when the subject’s alertness level was low. As a result, one run and two runs were discarded from subjects S1 and S2, respectively. Second, runs with large head motion (maximum translation ≥ 2 mm) were excluded. In this procedure, we excluded two runs from subject S5. After preprocessing the MRI data, we obtained 18–20 single-trial samples for each image and subject.

### Brain regions of interest

According to standard retinotopy experiments (Engel, et al., 1994; Sereno et al., 1995), we delineated V1, V2, V3, and V4. The lateral occipital complex (LOC), fusiform face area (FFA), and parahippocampal place area (PPA) were identified using conventional functional localizers (Kanwisher et al., 1997; Epstein and Kanwisher, 1998; Kourtzi and Kanwisher, 2000). We defined the higher visual cortex (HVC) region by manually delineating a contiguous region that covered the LOC, FFA, and PPA on the flattened cortical surfaces. The visual cortex (VC) was defined by combining V1–V4 and the HVC.

### DNN image features

We defined the unit activations of a deep neural network (DNN) with visual image inputs as stimulus features. For the DNN, we used a variant of AlexNet, BAIR/BVLC CaffeNet model (Krizhevsky, et al., 2012) pretrained with images in ImageNet to classify 1000 object categories (the pre-trained model is available from https://github.com/BVLC/caffe/tree/master/models/bvlc_reference_caffenet). The CaffeNet model has five convolutional layers and three fully connected layers. We resized all stimuli to 227 *×* 227 pixels before feeding them into the CaffeNet model. We reshaped the outputs of each of the first seven layers (conv1–5, fc6, and fc7 layers; after the rectification operation, if not otherwise stated) to a vector for each visual image. The number of units in each of the CaffeNet layers is as follows: conv1, 209, 400; conv2, 186, 624; conv3 and conv4, 64, 896; conv5, 43, 264; fc6 and fc7, 4, 096.

### DNN feature decoding

We constructed multivoxel decoders by training a set of linear regression models that predicted stimulus features from multiple fMRI voxel signals induced by the corresponding stimuli, as in previous studies(Horikawa and Kamitani, 2017; Shen et al., 2019a; Horikawa and Kamitani, 2022). Using fMRI samples from the training session (training dataset; 16,000 trials for subjects S1–4 and 6,400 trials for subjects S5–7), we trained a decoder for each combination of DNN units and brain areas (whole visual cortex or individual visual subareas). For a target DNN unit, we selected voxels that were most highly correlated (measured using absolute Pearson correlation coefficient) from each brain area using training data and then provided them to a decoder as inputs (with a maximum of 500 voxels). The weights of a decoder were optimized via least-square minimization with L2 regularization. We set the regularization parameter to 100.

We applied the trained decoders to fMRI data from the test dataset to predict the feature values of individual DNN units (“decoded features”). The decoded features corresponding to a visual image come from a single-trial fMRI sample. We normalized the decoded features for subsequent image reconstruction analyses to compensate for the possible differences in the distributions of the stimulus and decoded features. The variance across normalized decoded features within a layer was matched to the mean variance of DNN feature values, which was calculated from an independent set of 10,000 natural images. The mean of the normalized decoded features was maintained at the same level as that of the unnormalized decoded features.

### Image generator

#### GAN (main method)

To visualize the illusory percepts as images, we adopted a generator network (Dosovitskiy and Brox, 2016) that was pre-trained using a Generative Adversarial Network (GAN) framework (available from https://lmb.informatik.uni-freiburg.de/resources/binaries). This image generator was trained to transform the rectified outputs of fc6 layer of the CaffeNet model into the original input image using images from ImageNet (Russakovsky et al., 2015). We fed the decoded fc6 features from a single-trial or trial-averaged fMRI sample into this generator network to obtain a reconstructed image (VanRullen and Reddy, 2019). We chose this method for quantitative analyses in this study because the results of pilot research indicated that it tended to produce high-contrast reconstructions, especially for geometric shapes and patterns, compared to other candidate methods.

#### Diffusion

Similar to the main method, we fed the decoded fc6 features to a diffusion-based image generator. Here, we trained a conditional diffusion model after a modification of the model architecture from a previous study (Dhariwal and Nichol, 2021). The diffusion model was trained to generate the original input image conditioned on the rectified outputs of fc6 layer of the CaffeNet model as follows. We employed the architecture for the class-conditional ImageNet 64 *×* 64 model (available from https://github.com/openai/improved-diffusion/), which had a conditioning vector with a dimensionality of 768. To enable training a diffusion model conditioned on the 4096-dimensional fc6 feature vector derived from CaffeNet (see “DNN image features”), we further added a fully-connected layer whose input and output sizes were 4,096 and 768, respectively. The fc6 features were fed into the diffusion model through this fully-connected layer. Approximately 1.2 million natural images from ImageNet(Russakovsky et al., 2015) were used as training images and resized to 6464 pixels. We used the linear noise schedule and set the number of diffusion steps to 4000. The model was trained for 1 million training steps using a batch size of 128 and a learning rate of 0.0001.

#### Pixel optimization (iCNN)

This method was used to convert decoded features of multiple DNN layers to an image (code available from https://github.com/KamitaniLab/DeepImageReconstruction) in our original deep image reconstruction study (Shen, et al., 2019a). The pixel values of an input image were optimized such that its image features matched the decoded features. In the current study implementation, we used CaffeNet as the target of feature decoding for comparison with the other methods. Following Shen et al. (2019a), we used the feature values before the rectification operation from eight layers (conv1–5 and all fully connected layers). We also applied the same loss function and natural image prior, and solved the optimization problem using stochastic gradient descent with momentum for 200 iterations.

### Analysis of robustness to noise

To exclude the possibility that stimulus-independent noise in brain-decoded features leads to the reconstruction of illusory components, we added noise to stimulus fc6 features and fed them into the generator. Lacking prior knowledge of the noise distribution, we adopted a non-parametric method to sample the noise. We assumed that the noise for an individual DNN unit follows the same unknown probability distribution across the non-illusory trials of the same subject (individual units do not necessarily share the same distribution). The following analysis was performed separately for each subject. We calculated the empirical noise distributions by pooling the differences between decoded and stimulus features across non-illusory trials. We then randomly sampled the noise value from the empirical distribution for each DNN unit and added them to the stimulus features of an illusory image.

### Evaluation of reconstruction

#### Illusory line

We evaluated each single-trial reconstructed image of the line illusion, both globally (whole image) and locally (restricted to a specific image region). For the local case, we investigated two types of image regions: illusory and non-illusory. To delineate the regions for each stimulus image, we cropped the four largest disks tangent to the middle lines of the image, among which two disks consisted of illusory lines and two disks did not (Fig. 3E). We detected the principal orientation for each image/region of interest and compared which of the illusory and inducer orientations was more similar to the principal orientation, using cosine similarity. The principal orientation was detected using the method that has been used in texture analysis (Jafari-Khouzani and Soltanian-Zadeh, 2005). We converted the images into grayscale and applied Radon transform to detect the linear trends. More specifically, the largest disk area A in the image region was selected and projected it to a line space by summing the pixel intensities along each line within A:

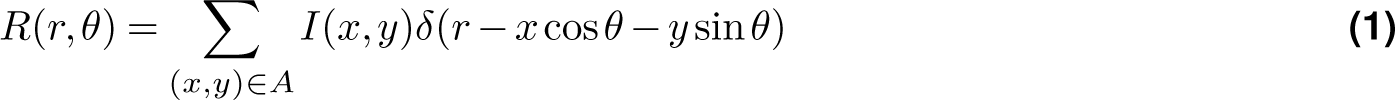

where each line is parametrized by the distance from the center *r* and the orientation *θ*. The intensity of the pixel located at (*x, y*) is denoted by *I*(*x, y*). If (*x, y*) is on the line, *δ* = 1; otherwise, *δ* = 0.

For each orientation, we calculated the variance of the projections across lines that intersect a small disk region in the center of the image. Intuitively, a prominent black line at a specific orientation would result in a significant decrease in the projection value, causing a sharp change in projections across the neighboring lines of the same orientation. We calculated this variance for different orientations and defined the principal orientation as the one with the largest variance. Only the lines close (no more than five pixels) to the center of the region were used to calculate variance. This restricted range was expected to contain the illusory line.

#### Illusory color

To evaluate the color filling-in effect, we performed a regression analysis in which a reconstructed image was approximated by the superposition of the illusory surface and the inducing stimulus with an additive error term. Given the real color surface in the positive control image, we performed another regression analysis that excluded the inducing stimulus regressor to compare the illusory color with the real color.

Before the regression analysis, we created redness maps from the reconstructions, the illusory/real surfaces, and the inducing stimulus. A redness map was generated by converting the RGB image to the HSV color space, using version 4.5.2 of the openCV library, and by extracting the saturation (S) values of the red pixels. The non-red pixels were assigned a value of zero. The red pixels were identified as those with hue values (H) in the range of 0◦ to 10◦, or 160◦ to 180 ◦. The regression models for the redness map of each single-trial reconstruction were as follows:

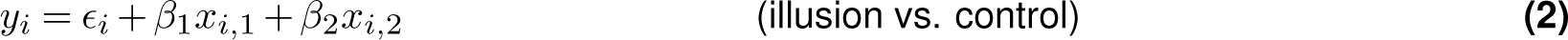

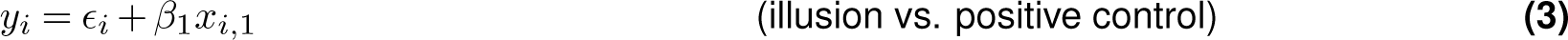

with *y_i_* (*i* = 1, 2, *… , h × w*) the value of the *i*-th pixel in the redness map of a reconstruction of size *h × w*, *E_i_* the error term, *x_i_*_,1_ and *x_i_*_,2_ the values of the *i*-th pixel in the redness maps of the illusory/real surface and the inducing stimulus, *β*_1_ and *β*_2_ the coefficients for the illusory/real surface and the inducing stimulus. The coefficients were estimated by minimizing the sum of squared errors across the pixels in each reconstruction.

### Analysis of individual units’ responses

#### Orientation selectivity

To examine whether individual DNN units could respond similarly to real and illusory line orientations, we introduced illusory images comprised of concentric curves broken at the center, along with their corresponding positive control images with a central real line (fig. S4A). The spacing between adjacent concentric curves was kept constant, and the phases of the concentric halves were randomly assigned from a range of 0 to 1 cycle. To balance the background curves, the images with flipped phases between two parts were paired, and the averaged activation was used to evaluate orientation selectivity.

We first identified units selective to a specific orientation using a real central line of 12 different orientations (in 15◦ steps) on unbroken concentric patterns with phases matched with the illusion/positive control image. The concentric patterns were presented in the background because unit activation to a central orientation depended on the background context, especially in the middle and higher layers. Among center-responsive units (the center units of each channel in convolutional layers [864 for conv1, 256 for conv2 and conv5, 384 for conv3 and conv4] and all units in fully connected layers), we selected the units that exhibited higher activation to the orientation of interest than to all the other orientations for both of the background phases. We then ranked them based on the difference in activation between the orientation of interest and the other orientations. Using the top 5% units of each layer, we calculated the activations to all 12 orientations for the illusion and the positive control images. We repeated this procedure for 50 randomly assigned pairs of background phases.

#### Color (hue) selectivity

To examine whether individual DNN units could respond similarly to real and illusory colors, we identified units responsive to real color (fig. S5A) and then compared their activations to the illusion, the control, and the positive control images (fig. S5B; see “Visual stimuli”).

We first identified units selective to red color, using a uniform surface (disk or square) with three levels of luminance (0.3, 0.5, and 1, relative to the luminance of the image background color based on measurements of the display) and four levels of saturation (0, 0.3, 0.7, 1) on the inducer patterns. Among the units that showed higher activations to red (non-zero saturation) than to gray (zero saturation) for all the luminance and saturation levels, we ranked the units by the difference in activation averaged across all comparisons between the red and gray surfaces. The activations for the top 5% units are shown for each layer after subtracting the activation to the gray surface.

### Statistical tests and sample size

In this study, each subject was regarded as a replication unit (Ince, et al., 2022); thus, statistical tests were primarily conducted on a per subject basis. The sample size of test data (the number of trials for each image/condition) was determined before the test experiments.

In the anlaysis of principal orientations, we planned to perform one-sided z-tests to determine whether the closer-to-illusory proportion was greater in one condition than in the other. To achieve a statistical power of 80%, at least 20 samples are required for each condition to detect a large effect size (Cohen’s *h* = 0.8) at a significance level of 0.05. Thus, we repeated 20 trials for each stimulus image in each subject. In the present paper, we only present the results of statistical tests on the trials pooled across different stimulus configurations. Thus, one condition for statistical comparison involve more than 20 trials. The numbers of trials that survived the exclusion criteria for the results in Figure 3C to compare decoded features (*n*_1_) and stimulus features (*n*_2_) were *n*_1_ = *n*_2_ = 117, 116, 120, 120, 113, 120, 120 for subjects S1–7, respectively; in Figure 3D and F to compare local illusory (*n*_1_) and non-illusory (*n*_2_) regions,*n*_1_ = *n*_2_ = 234, 232, 240, 240, 226, 240, 240 for subjects S1–7.

In the analysis of illusory color, we planned to perform one-sided t-tests on the illusory surface coefficients obtained from individual trials/reconstructions. To achieve a statistical power of 80%, at least 20 samples are required for each condition to detect a large effect size (Cohen’s *d* = 0.8) at a significance level of 0.05. Thus, we repeated 20 trials for each stimulus image in each subject. The actual numbers of trials that survived the exclusion criteria in Figure 4E and F to compare the illusion (*n*_1_) and the control (*n*_2_) conditions were *n*_1_ = 77, 74, 80, 80, 75, 80, 80 and *n*_2_ = 79, 75, 80, 80, 77, 80, 80 [subjects S1–7] for Ehrenstein; *n*_1_ = 20, 20, 20,19, 20, 20 and *n*_2_ = 20, 20, 20, 19, 20, 20 [subjects S1–3 and S5–7] for Varin. The same numbers of trials/reconstructions were used for the results in Figure 4I and J.

**Figure S1.**
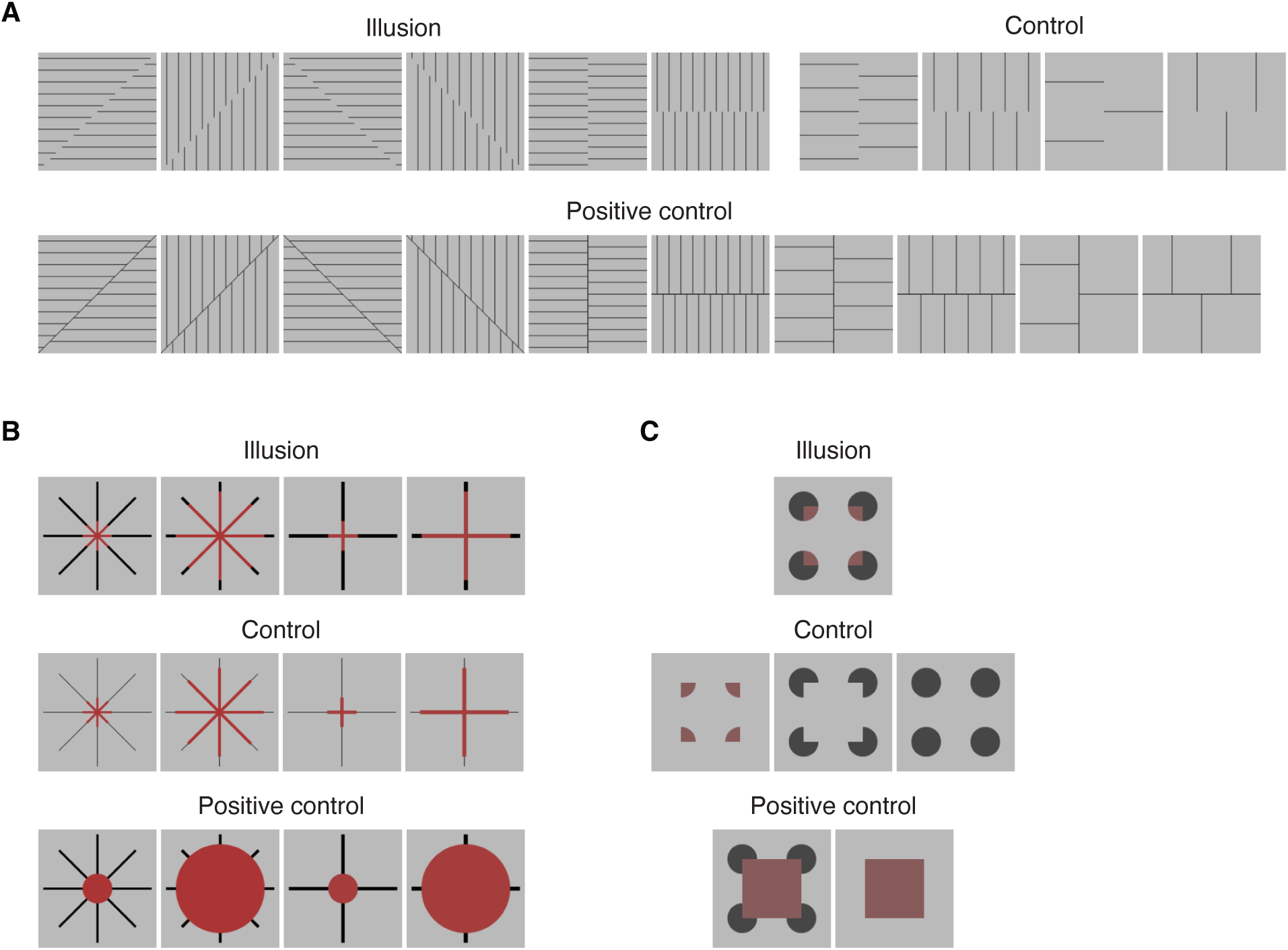
Test images used in fMRI experiment. (**A**) Example illusory and positive control images with the central line of different orientations. (**B**) Ehrenstein configuration for neon color spreading. (**C**) Varin configuration for neon color spreading.

**Figure S2.**
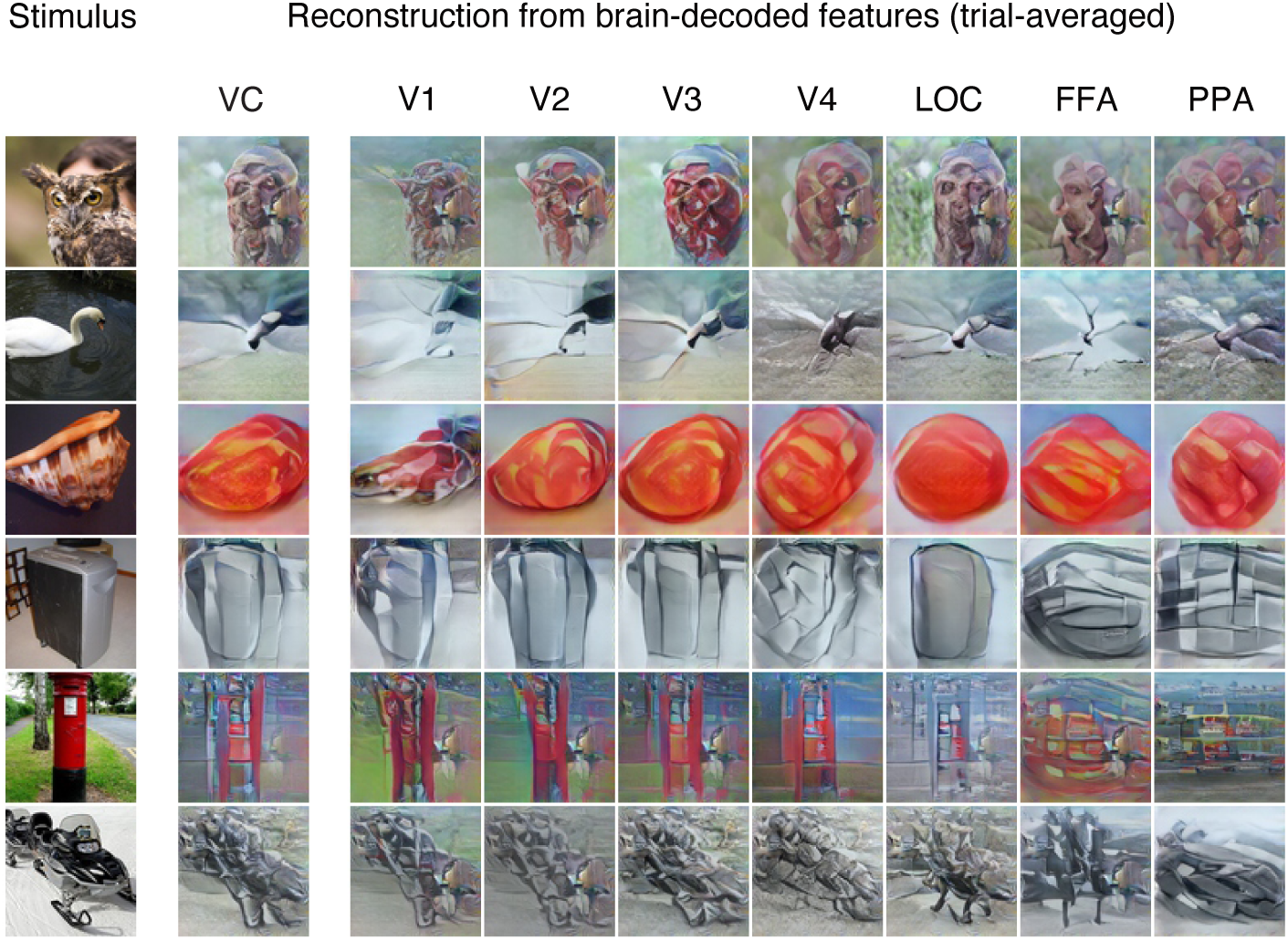
Reconstructions of natural images from brain-decoded features. The results of each column were produced from averaged fMRI signals across 24 trials in the whole visual cortex (VC) and individual visual areas of subject S2.

**Figure S3.**
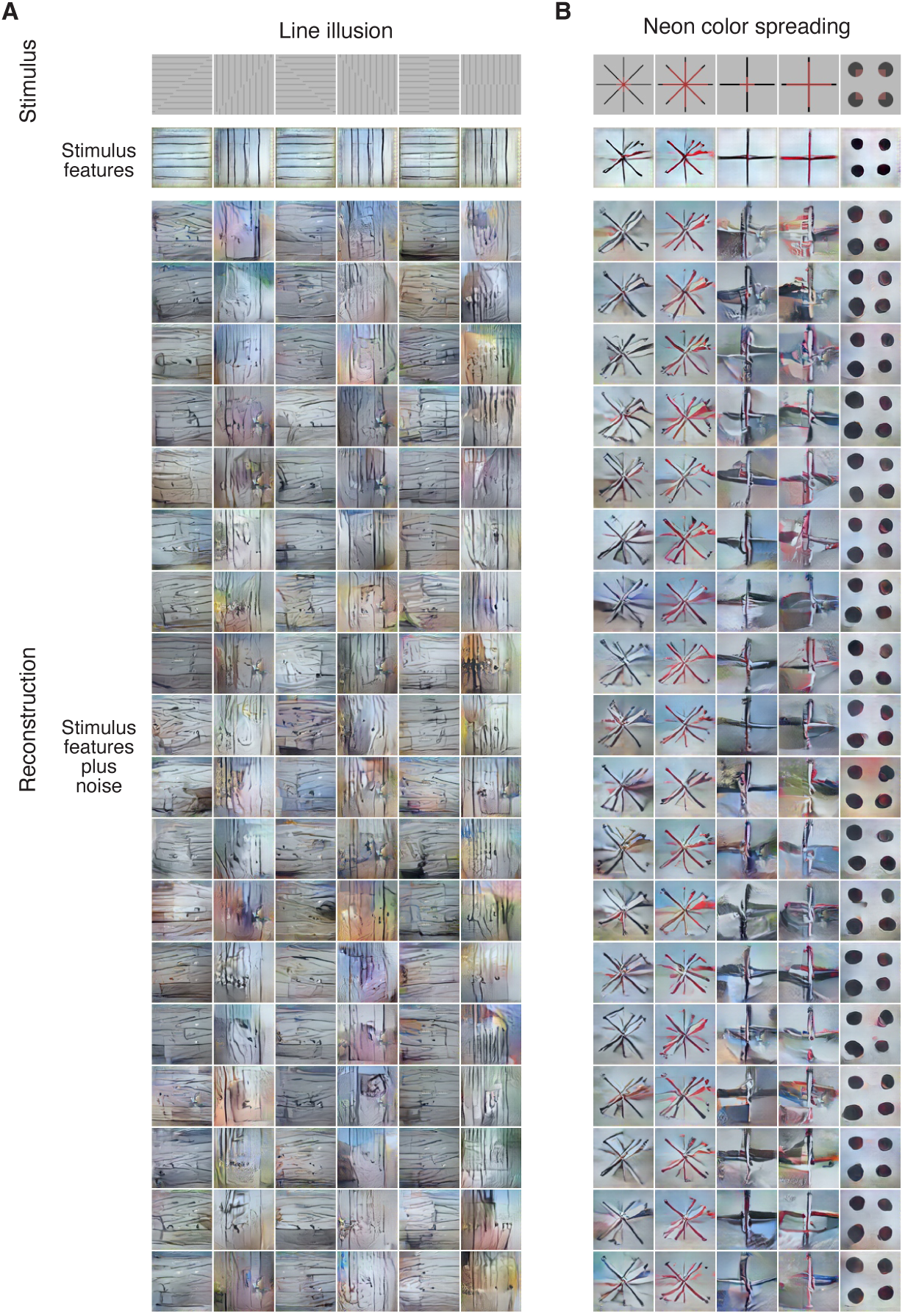
Reconstructions of illusory images from stimulus features plus the noise. The noises were sampled from the empirical noise distribution computed from the brain-decoded features of non-illusory images. (**A**) Line illusion.(**B**) Neon color spreading.

**Figure S4.**
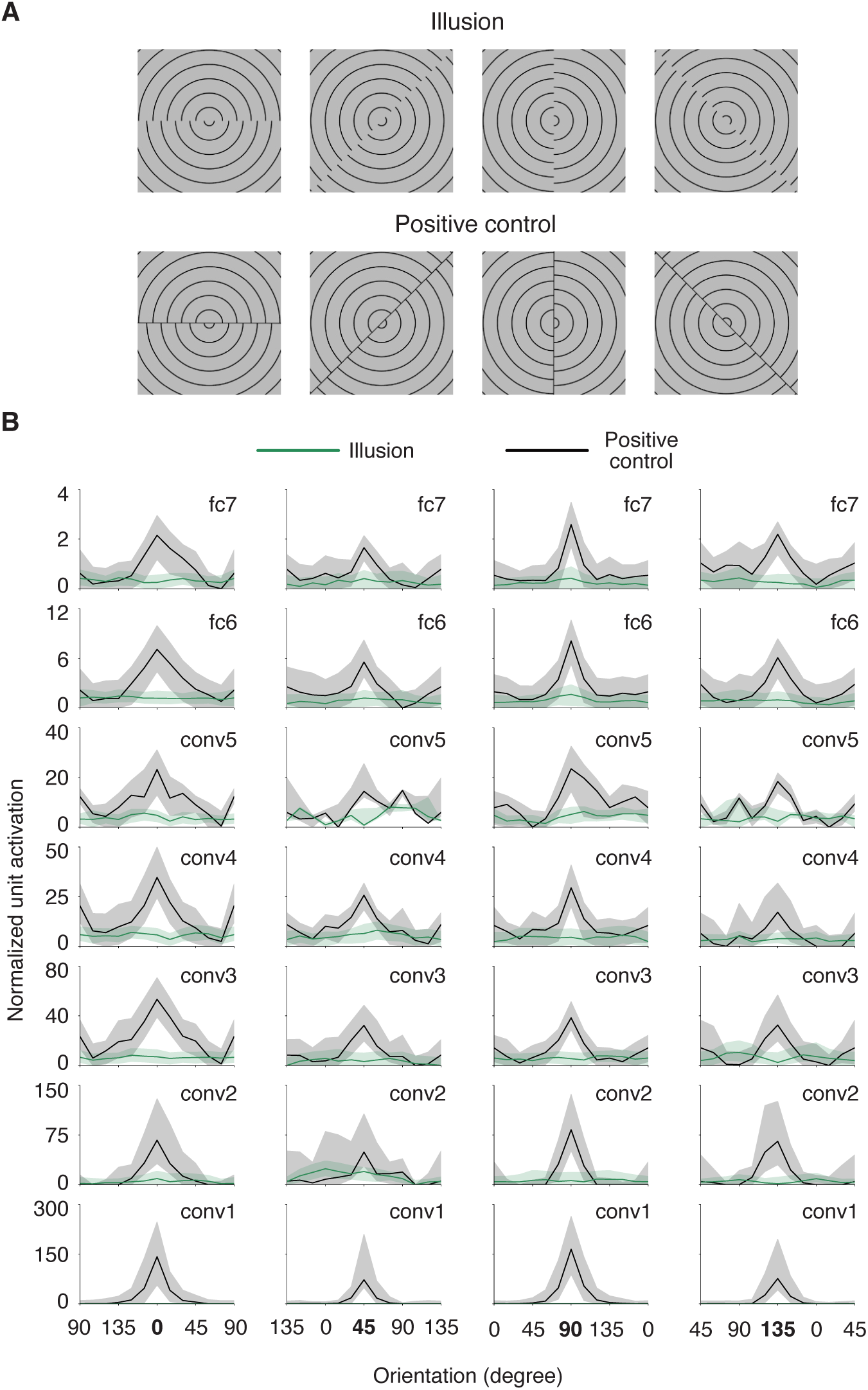
Individual DNN units do not show orientation tuning shared between illusory and real line. (**A**) Illusory line induced by offset-gratings. (**B**) Tuning curves of the orientation-selective units. The tuning curve of each unit was normalized by subtracting the minimum activation value of all orientations. Lines represent the median activation and shaded areas represent the interquartile range of the units pooled across different background phases. The tuning curves showed sharp peaks in the units’ preferred orientation for the positive control images (black) but not for the illusory images (green). The robustness of the results was confirmed when the central line of the positive control was white.

**Figure S5.**
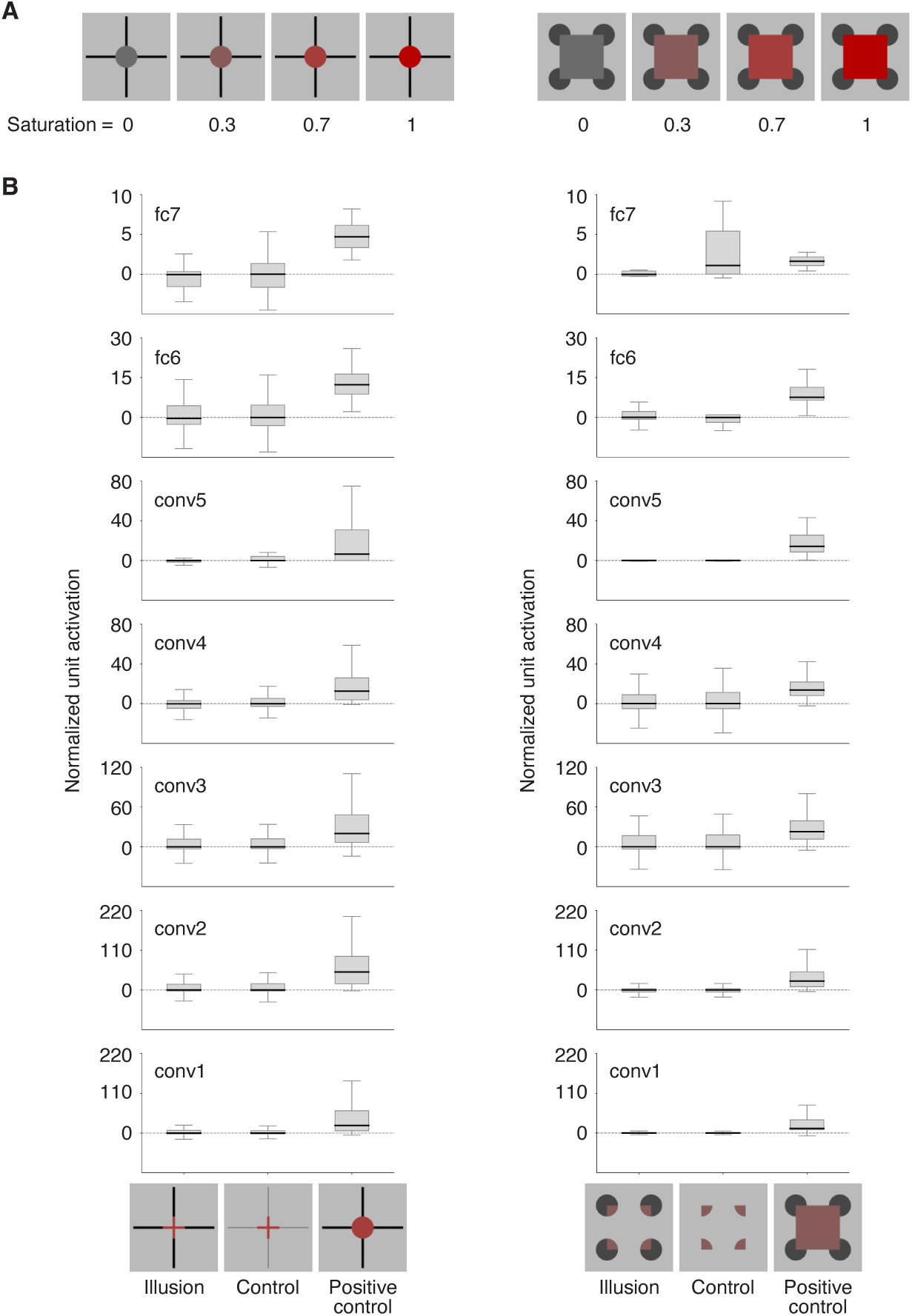
Individual DNN units do not respond to illusory color similar to real color. Units were analyzed for Ehrenstein (left) and Varin (right), respectively. (**A**) Example images used to identify red-sensitive units. (**B**) Normalized activation of color-selective units. Black lines represent the median value and shaded areas represent the interquartile range of units. If units respond to illusory color, the plots should show similar values for illusion and positive control images, which are larger than the control condition. However, illusion and control are almost identical, supporting the idea that units do not respond to illusory color. This small difference was not due to the lack of sensitivity of the units. In most units, control and illusion showed identical activation, or control showed relatively higher activation, not the other way around.

**Figure S6.**
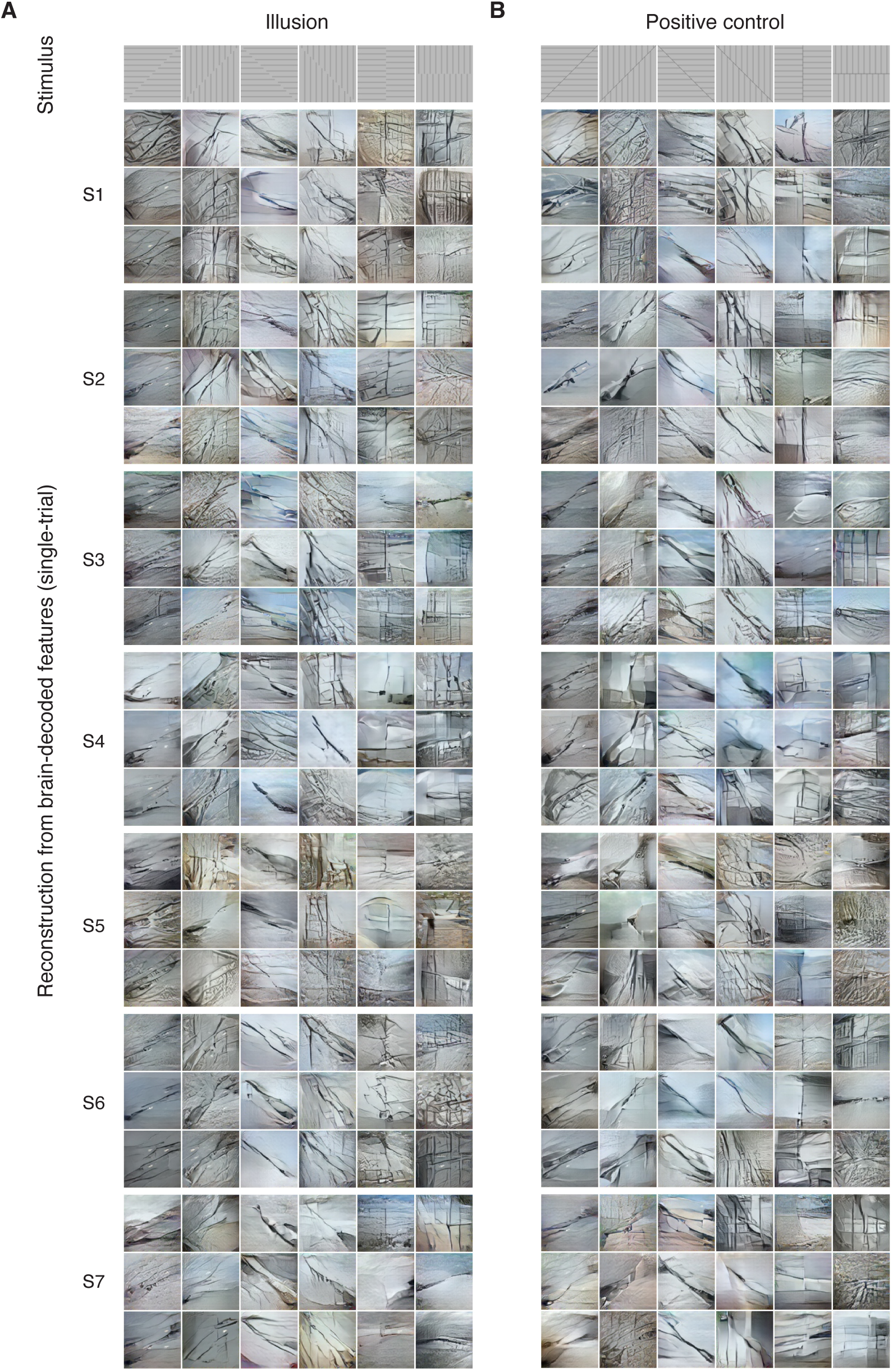
Reconstructions of line illusion for different configurations from brain-decoded features. Results were produced from single-trial fMRI signals in the whole visual cortex (VC). Representative reconstructions from three different trials are shown from each subject (no overlapping trials with Figure 2). (**A**) Illusion condition.(**B**) Positive control condition.

**Figure S7.**
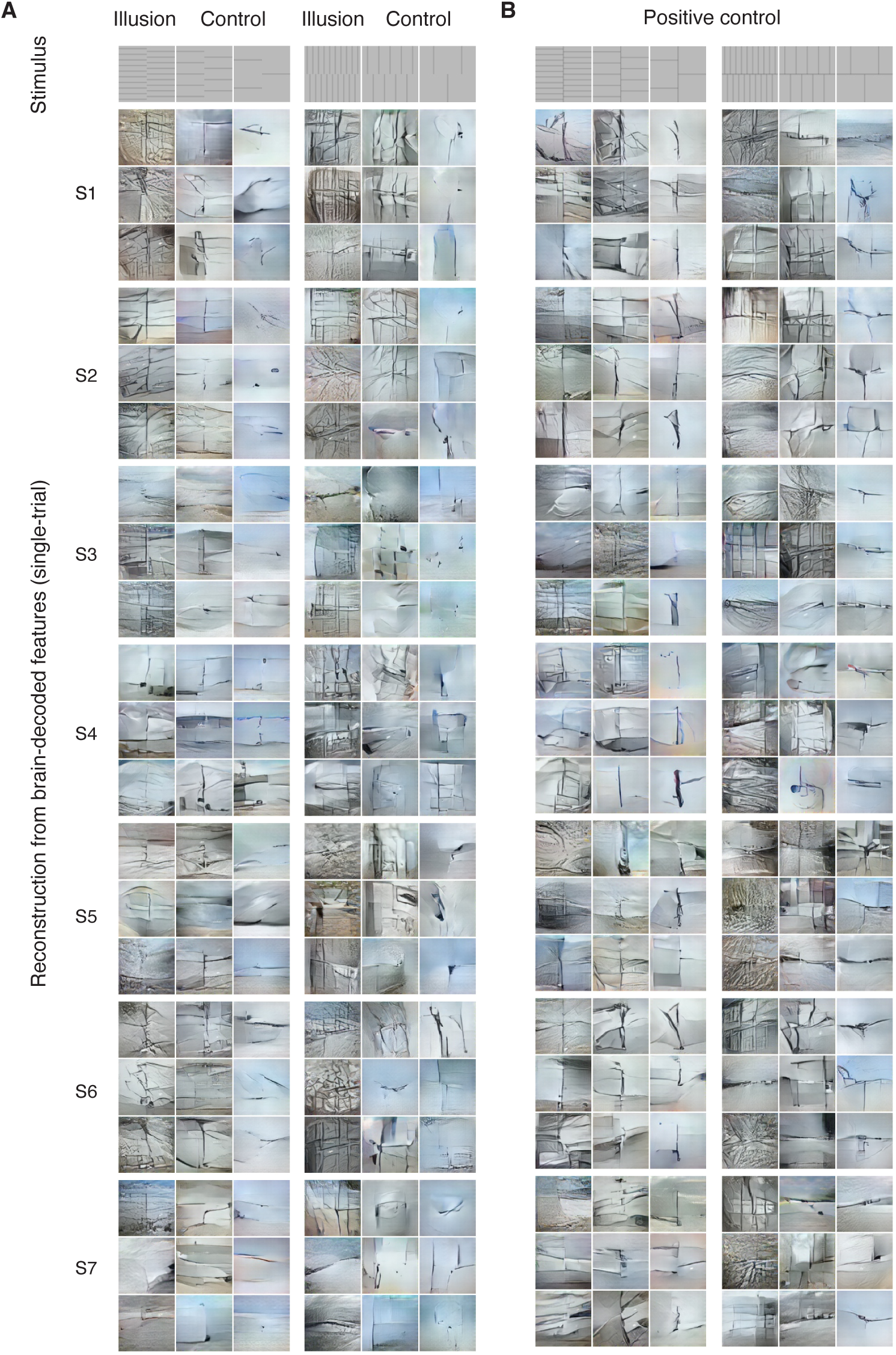
Reconstructions of illusory and control images from brain-decoded features for line illusion. Results were produced from single-trial fMRI signals in the whole visual cortex (VC). Representative reconstructions from three different trials are shown from each subject (no overlapping trials with Figure 2). (**A**) Illusion and control condition.(**B**) Positive control condition.

**Figure S8.**
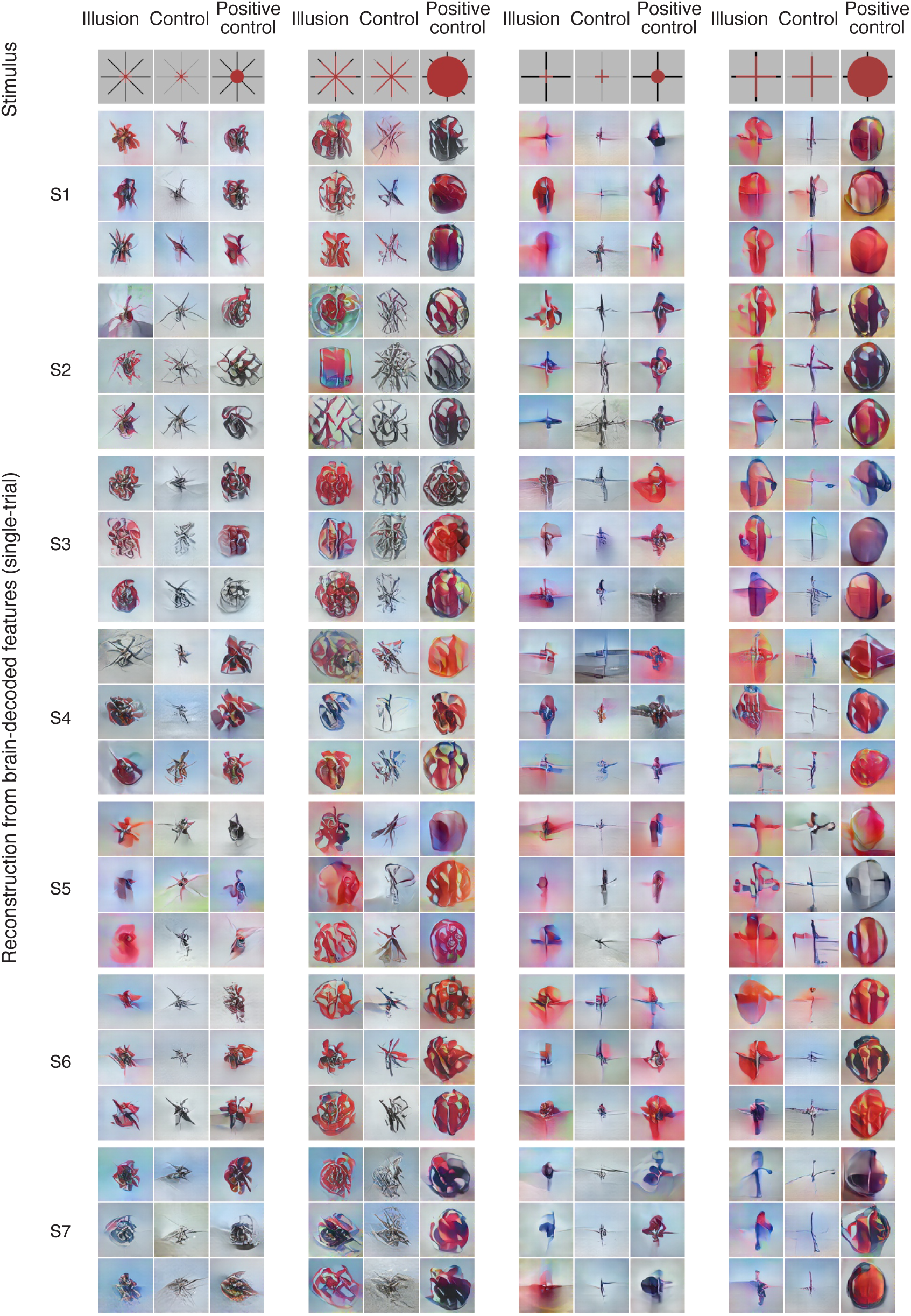
Reconstructions of neon color spreading (Ehrenstein) from brain-decoded features. Results were produced from single-trial fMRI signals in the whole visual cortex (VC). Representative reconstructions from three different trials are shown from each subject (no overlapping trials with Figure 2). Every three columns show illusion (left), control (middle), and positive control (right) conditions of the same configuration.

**Figure S9.**
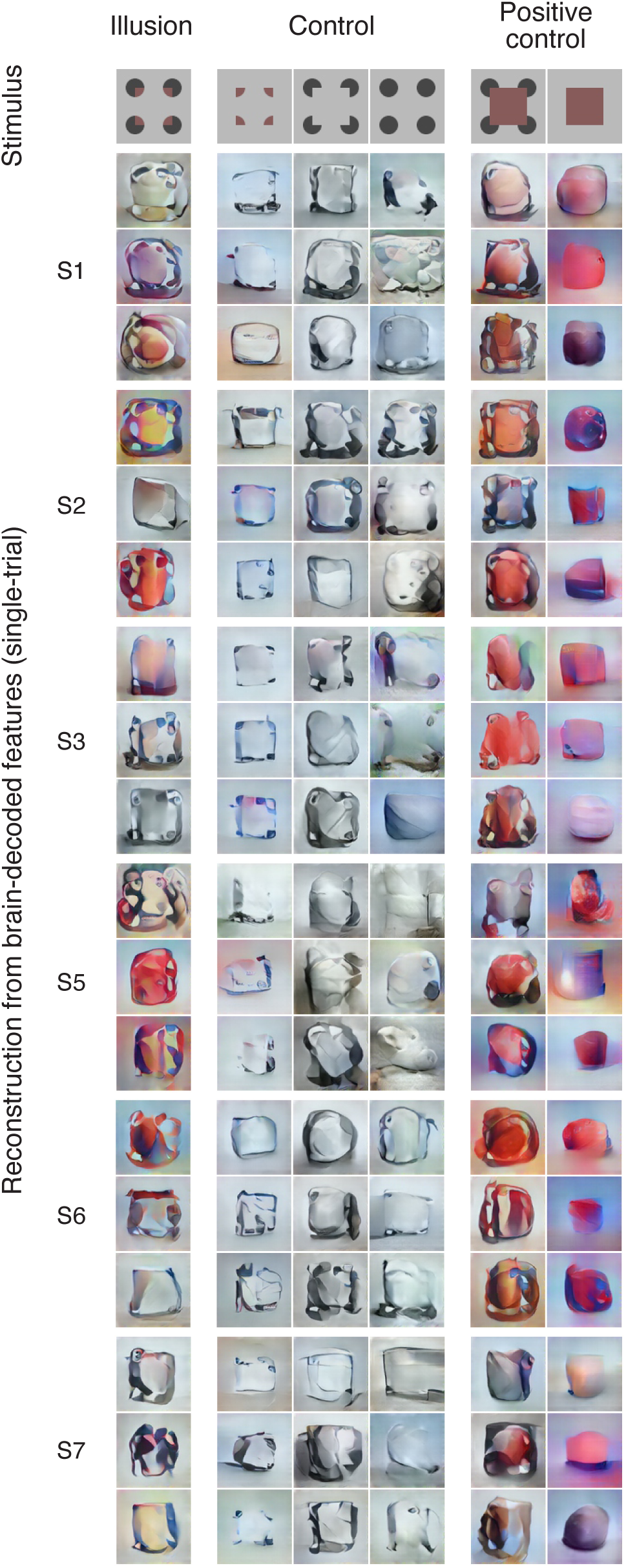
Reconstructions of neon color spreading (Varin) from brain-decoded features. Results were produced from single-trial fMRI signals in the whole visual cortex (VC). Representative reconstructions from three different trials are shown from each subject (no overlapping trials with Figure 2). The three panels show illusion (left), control (middle), and positive control (right) conditions.

**Figure S10.**
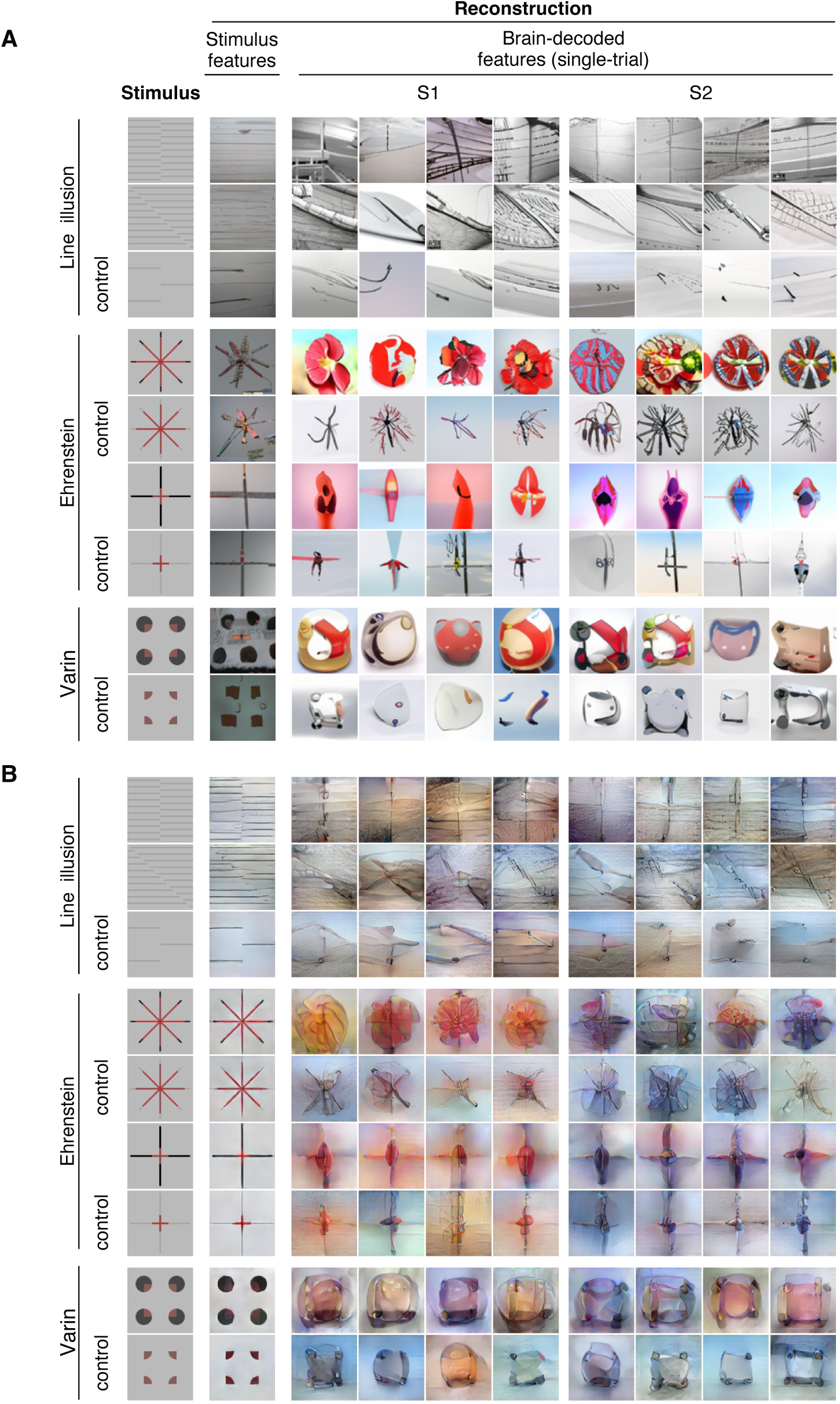
Reconstructions of illusory and control images using other generators. Reconstructions from stimulus features and from brain-decoded features are shown for two representative subjects (S1, S2). Reconstructions from brain-decoded features were produced from single-trial fMRI signals (same trials as those shown in Figure 2) in the whole visual cortex (VC). (**A**)Diffusion. (**B**) Pixel optimization (iCNN).

**Figure S11.**
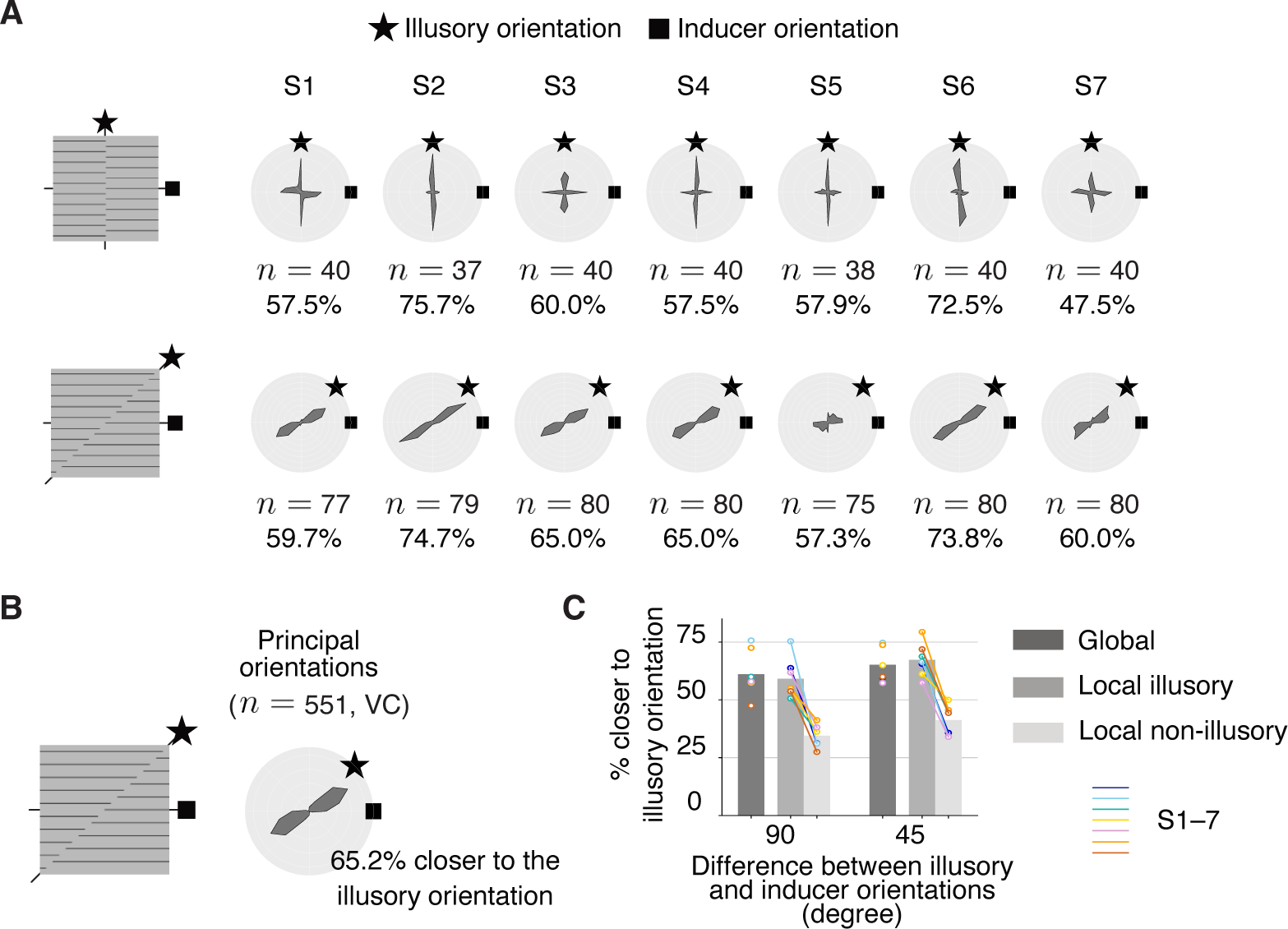
Evaluation of line illusion reconstructions for 90*^◦^*- and 45*^◦^*-difference configurations. Results are based on single-trial reconstructions from VC. (**A**)Distributions of principal orientations in reconstructions from individual subjects. The results pooled across 90*^◦^*- (top) and 45*^◦^*- (bottom) difference configurations are shown for each subject (totalling *n* samples; bin size = 15*^◦^*). (**B**) Distribution of principal orientations in reconstructions for 45*^◦^*-difference configurations (pooled across seven subjects, totalling *n* samples; bin size = 15*^◦^*). (**C**) Proportions of principal orientations closer to the illusory than the inducer orientation. Color circles and lines indicate individual subjects.

**Figure S12.**
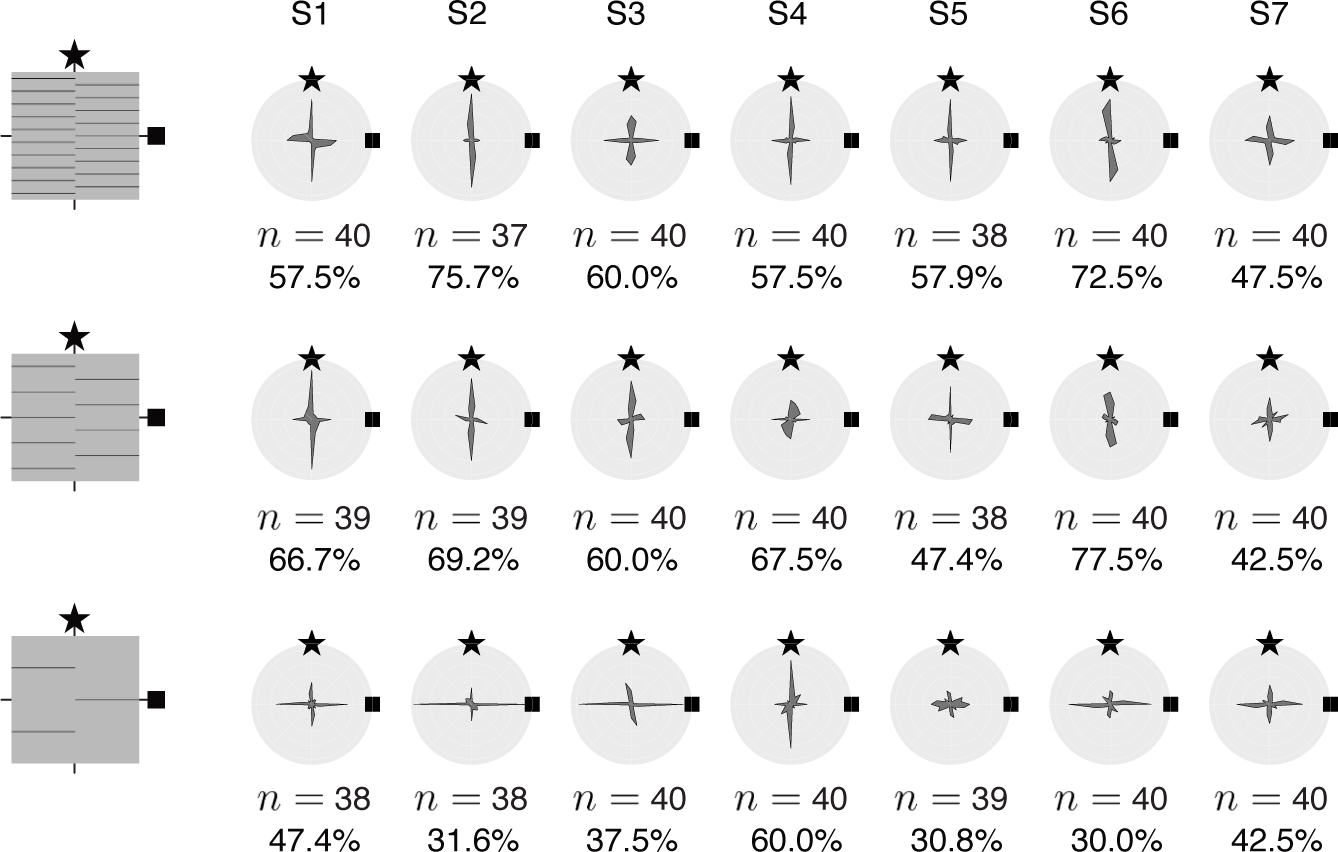
Evaluation of line illusion reconstructions for different numbers of inducer lines. Distributions of principal orientations in single-trial reconstructions from VC (results for 90*^◦^*-difference configurations are pooled for each subject, totalling n samples; bin size = 15*^◦^*).

**Figure S13.**
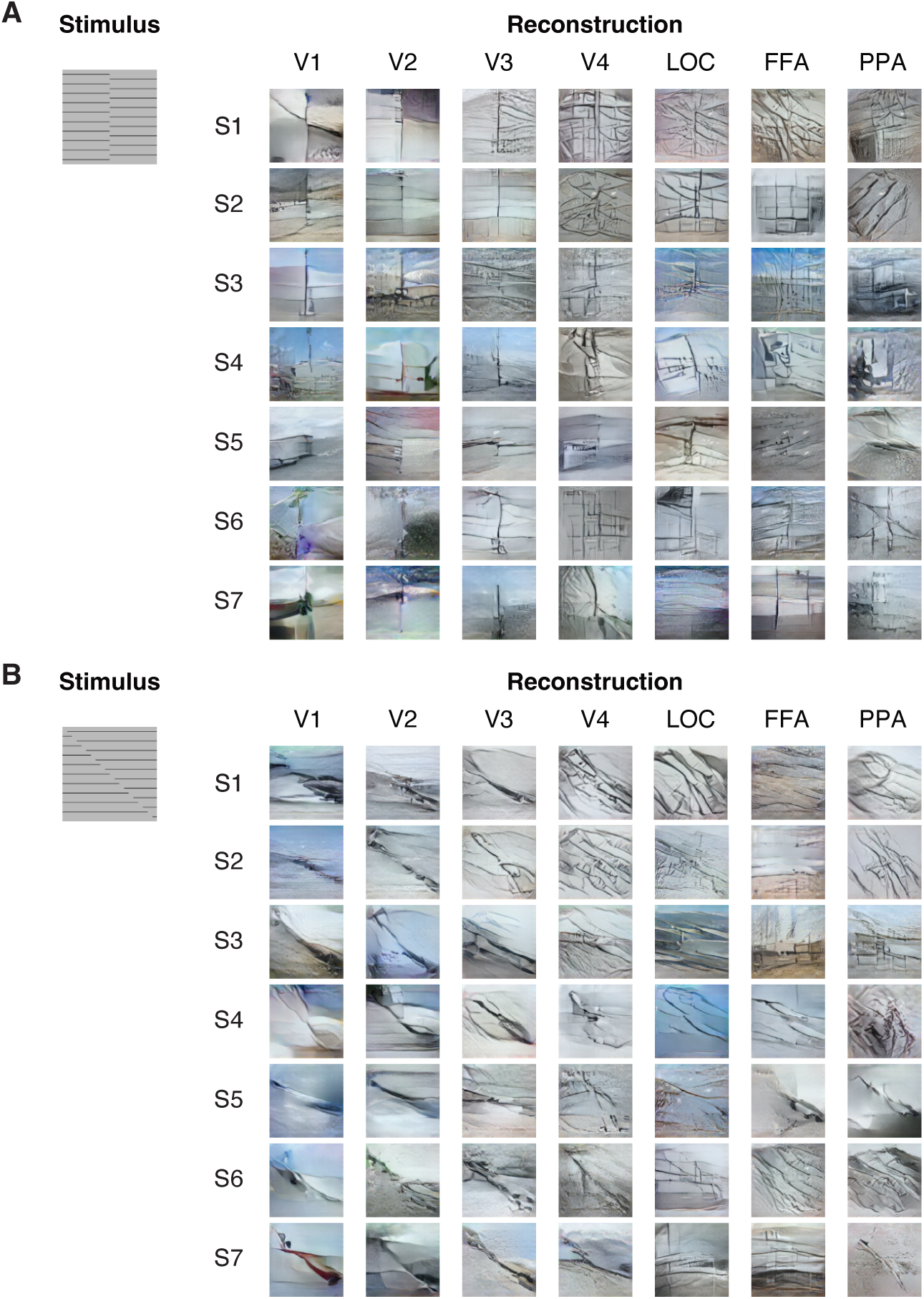
Reconstructions of line illusion from individual visual areas. Representative reconstructions from single-trial brain activity are shown for each subject (no overlapping trials with Figure 3). (**A**)90*^◦^*-difference configuration. (**B**) 45*^◦^*-difference configuration.

**Figure S14.**
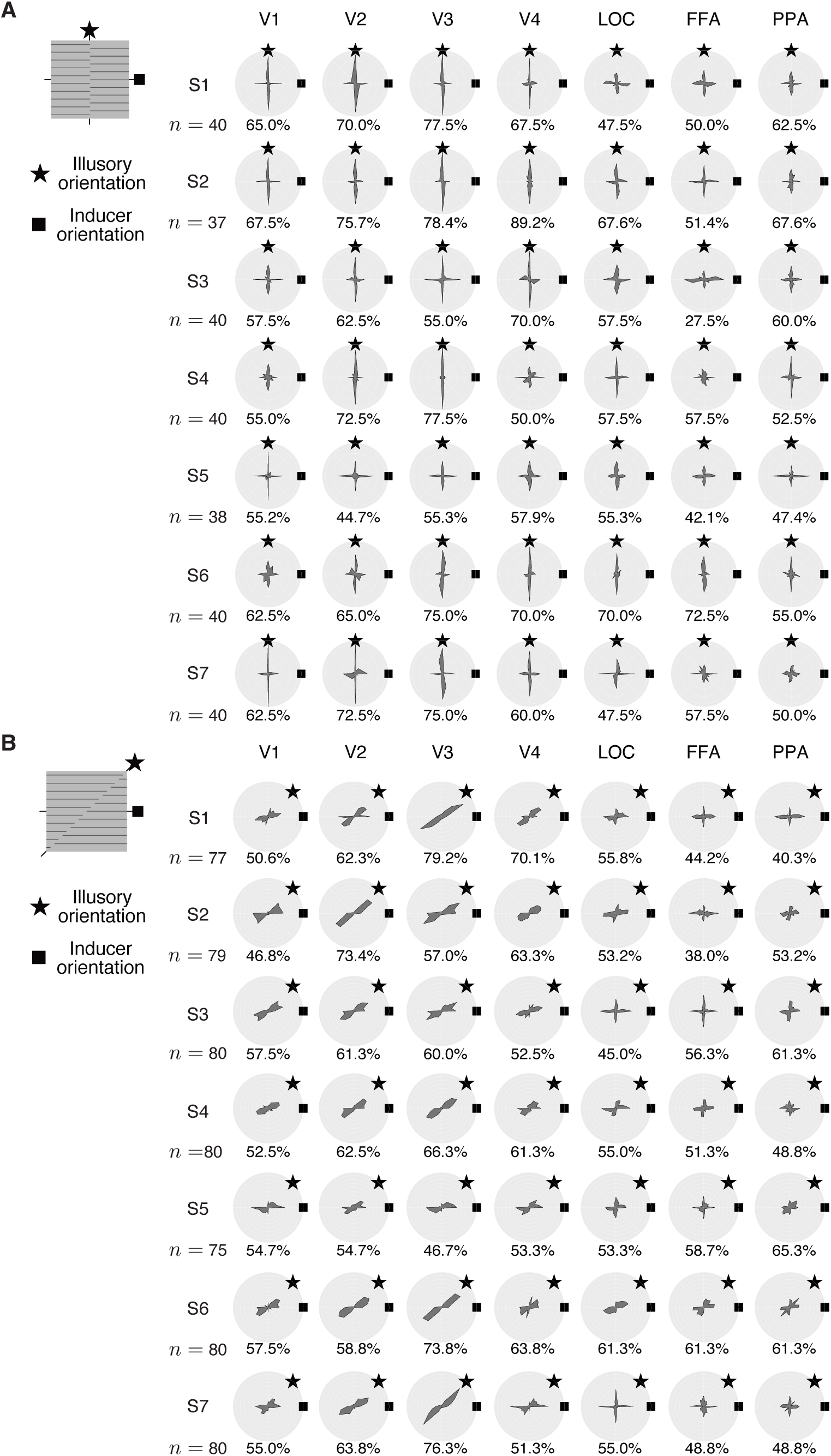
Evaluation of line illusion reconstructions from individual visual areas. Distributions of principal orientations in single-trial reconstructions are shown for each subject (pooled across all 90*^◦^*- or 45*^◦^*-difference configurations, totalling *n* samples; bin size = 15*^◦^*). (**A**)90*^◦^*-difference configuration. (**B**) 45*^◦^*-difference configuration.

**Figure S15.**
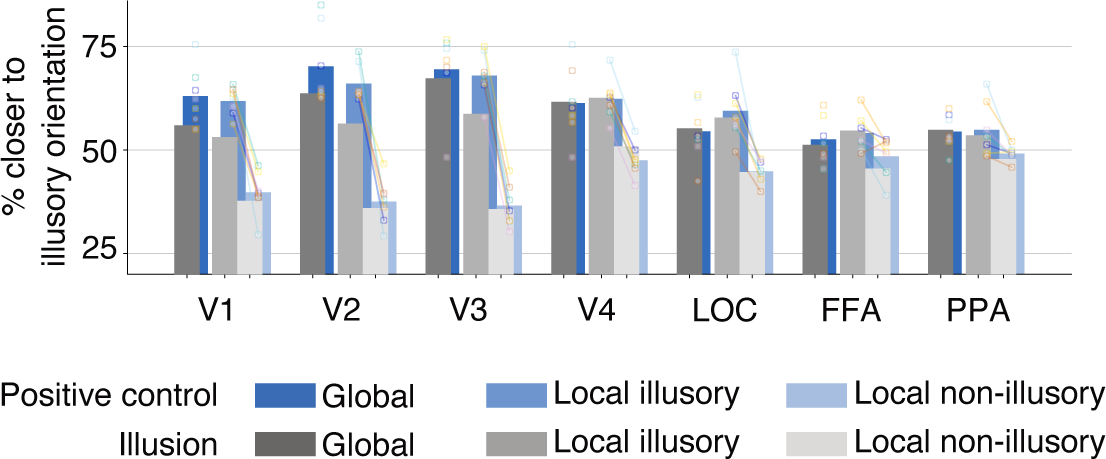
Comparison between positive control and illusion conditions for line illusion. Proportions of principal orientations closer to the illusory than to the inducer orientation are shown for individual visual areas (pooled across all subjects and configurations; gray bars are the same as that in Figure 3F). Color circles and lines indicate individual subjects for the positive control condition.

**Figure S16.**
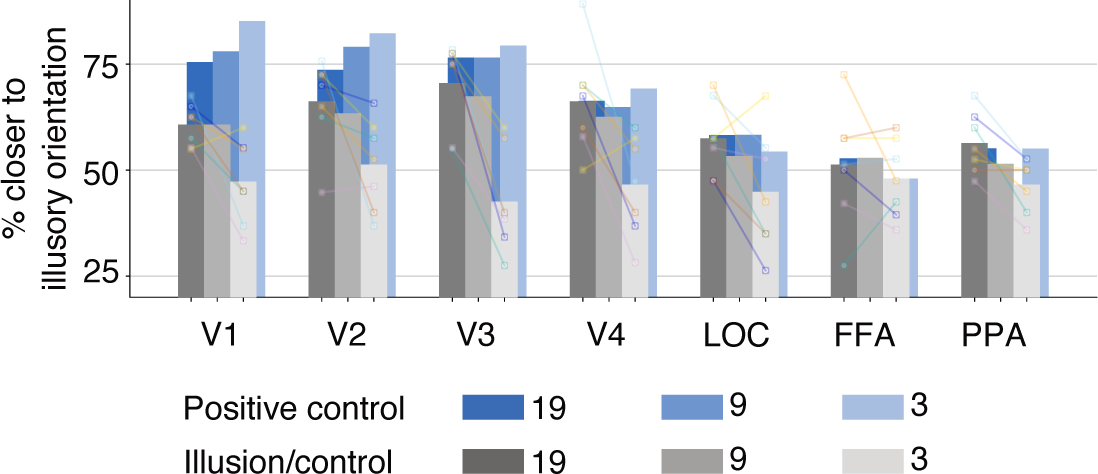
Comparison between different numbers of inducer lines for line illusion. Proportions of principal orientations closer to the illusory than to the inducer orientation in global regions are shown for individual visual areas. Color circles and lines indicate individual subjects for illusion or control conditions.

**Figure S17.**
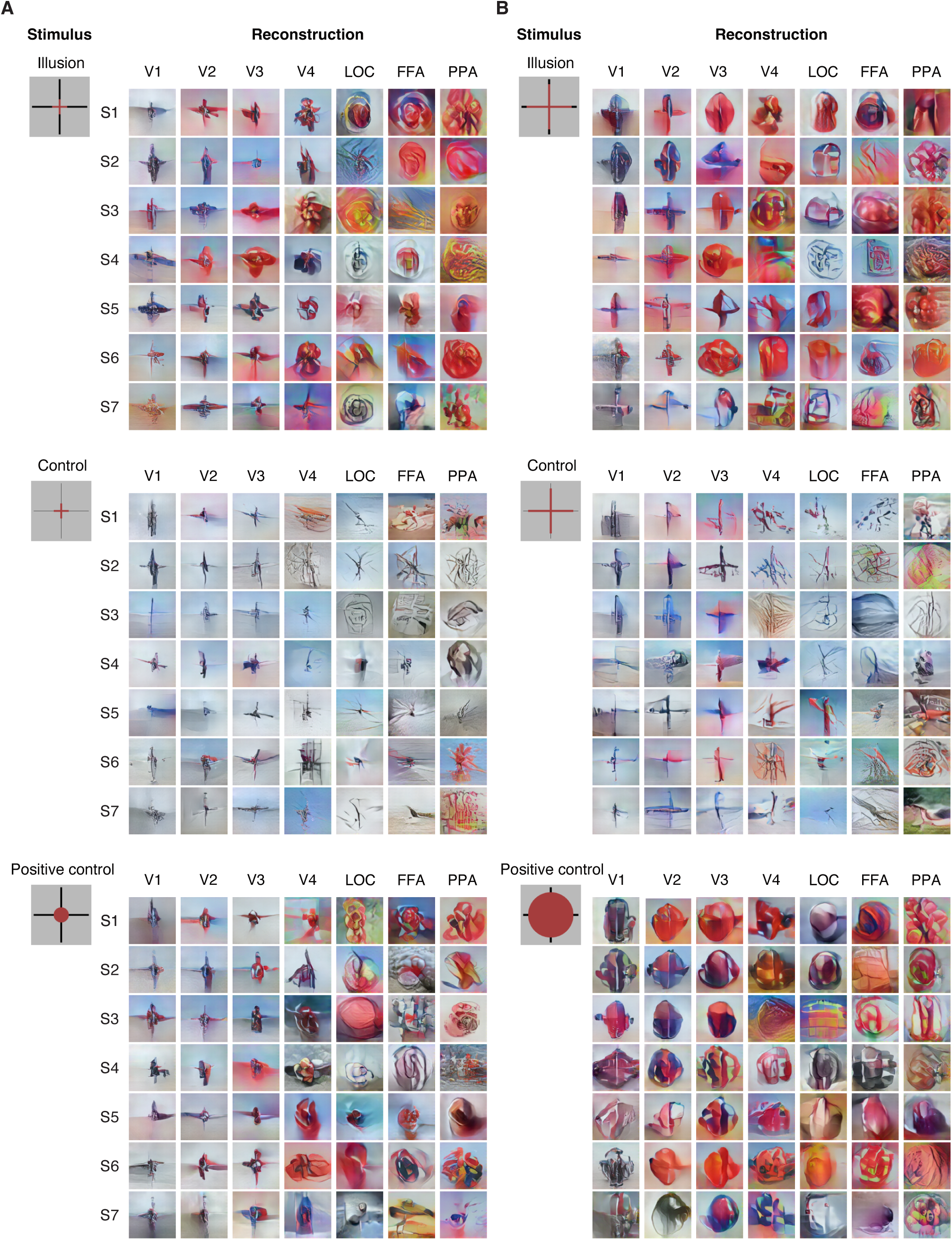
Reconstructions of neon color spreading (Ehrenstein) from individual visual areas. Representative single-trial reconstructions of the illusion (top), control (middle), and positive control (bottom) conditions are shown for each subject (no overlapping trials with Figure 4). (**A**) Small size. (**B**) Large size.

**Figure S18.**
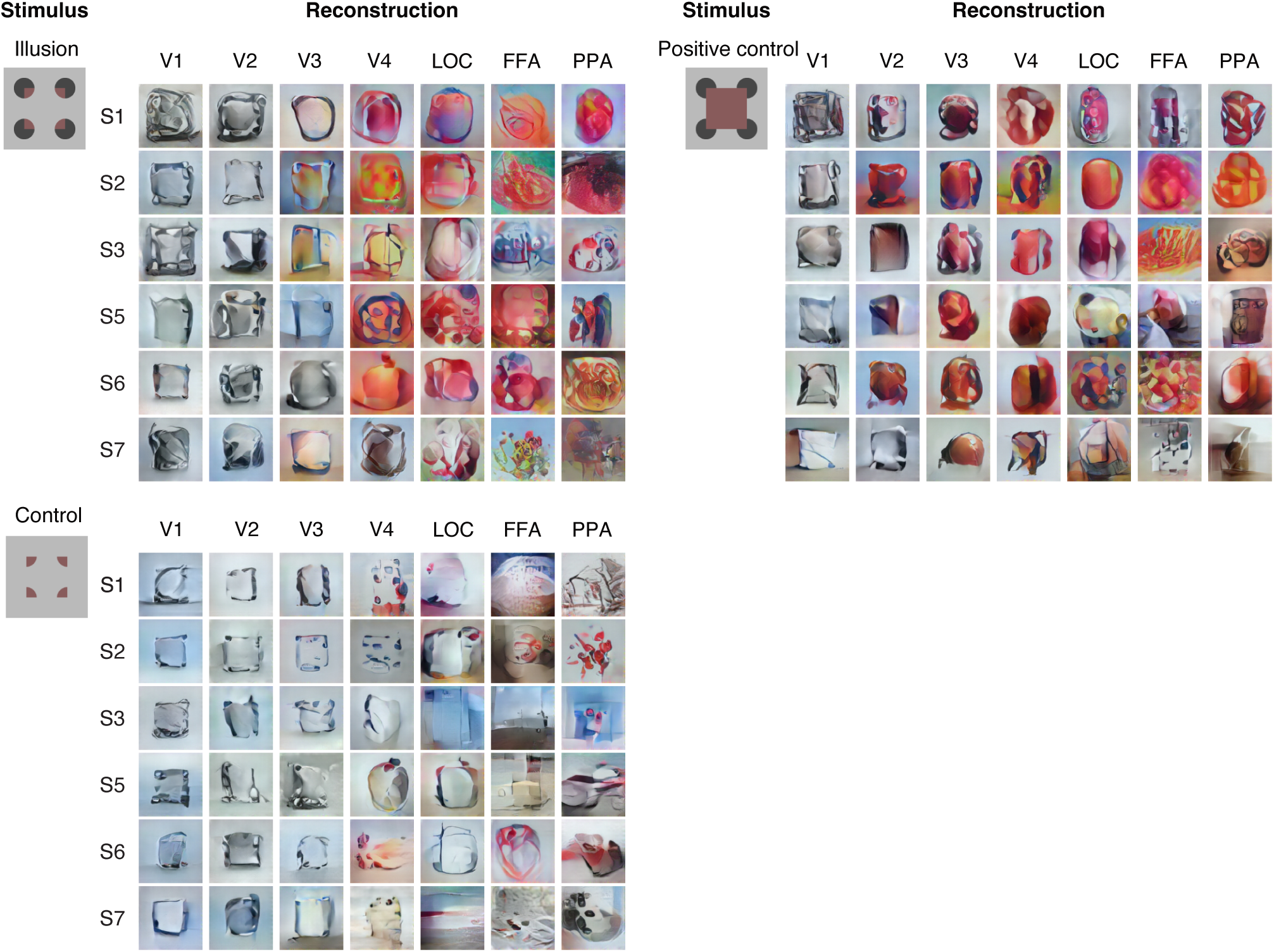
Reconstructions of neon color spreading (Varin) from individual visual areas. Representative single-trial reconstructions of the illusion (left top), control (left bottom), and positive control (right) conditions are shown for each subject (no overlapping trials with Figure 4).

**Figure S19.**
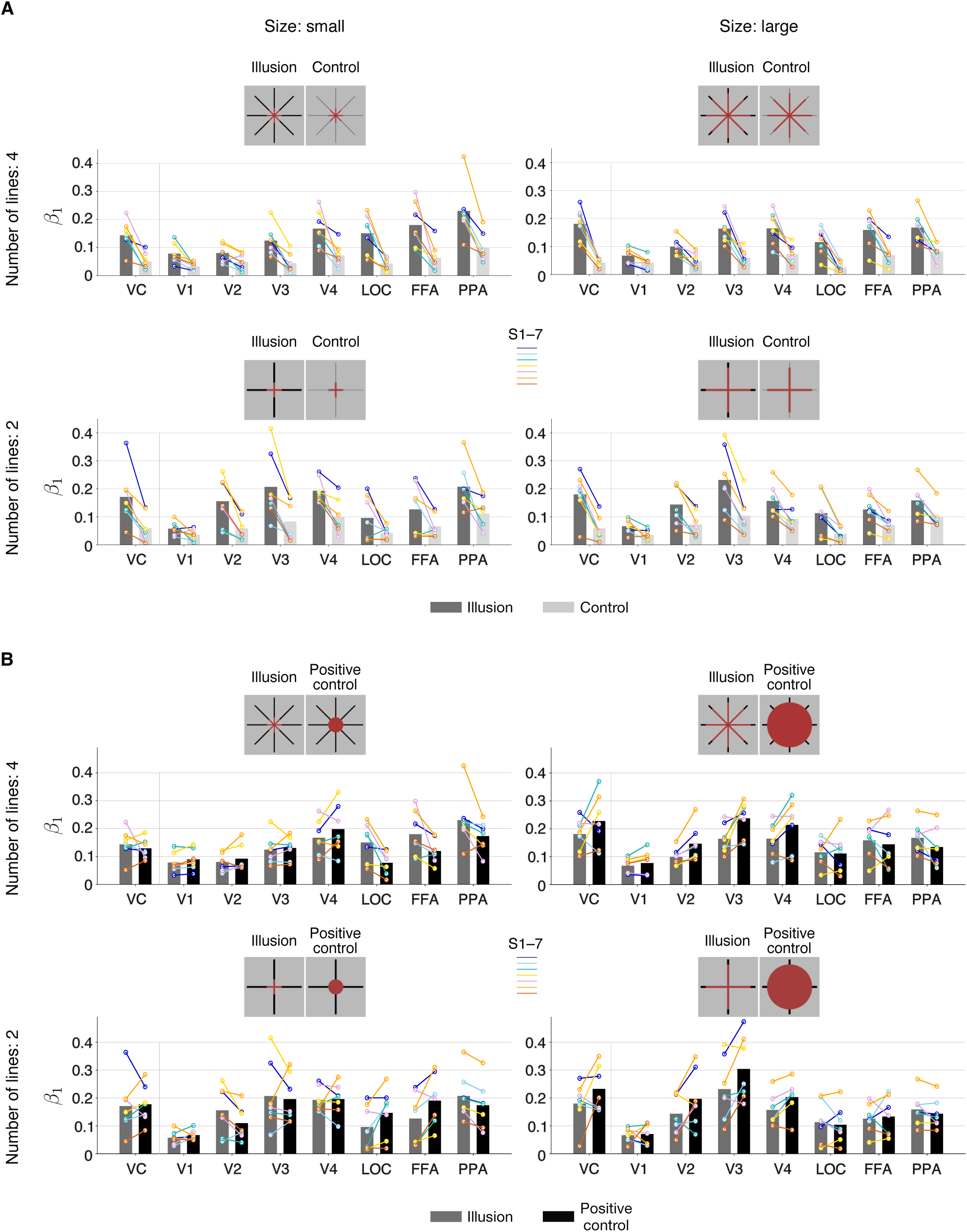
Evaluation of neon color spreading (Ehrenstein) reconstructions for different sizes and numbers of lines. Results are based on single-trial reconstructions from VC and individual visual areas. Color circles and lines indicate individual subjects. (**A**) Comparison of the illusory surface coefficient values between illusion and control conditions. (**B**) Comparison of the illusory surface coefficient values between illusion and positive control conditions.

**Figure S20.**
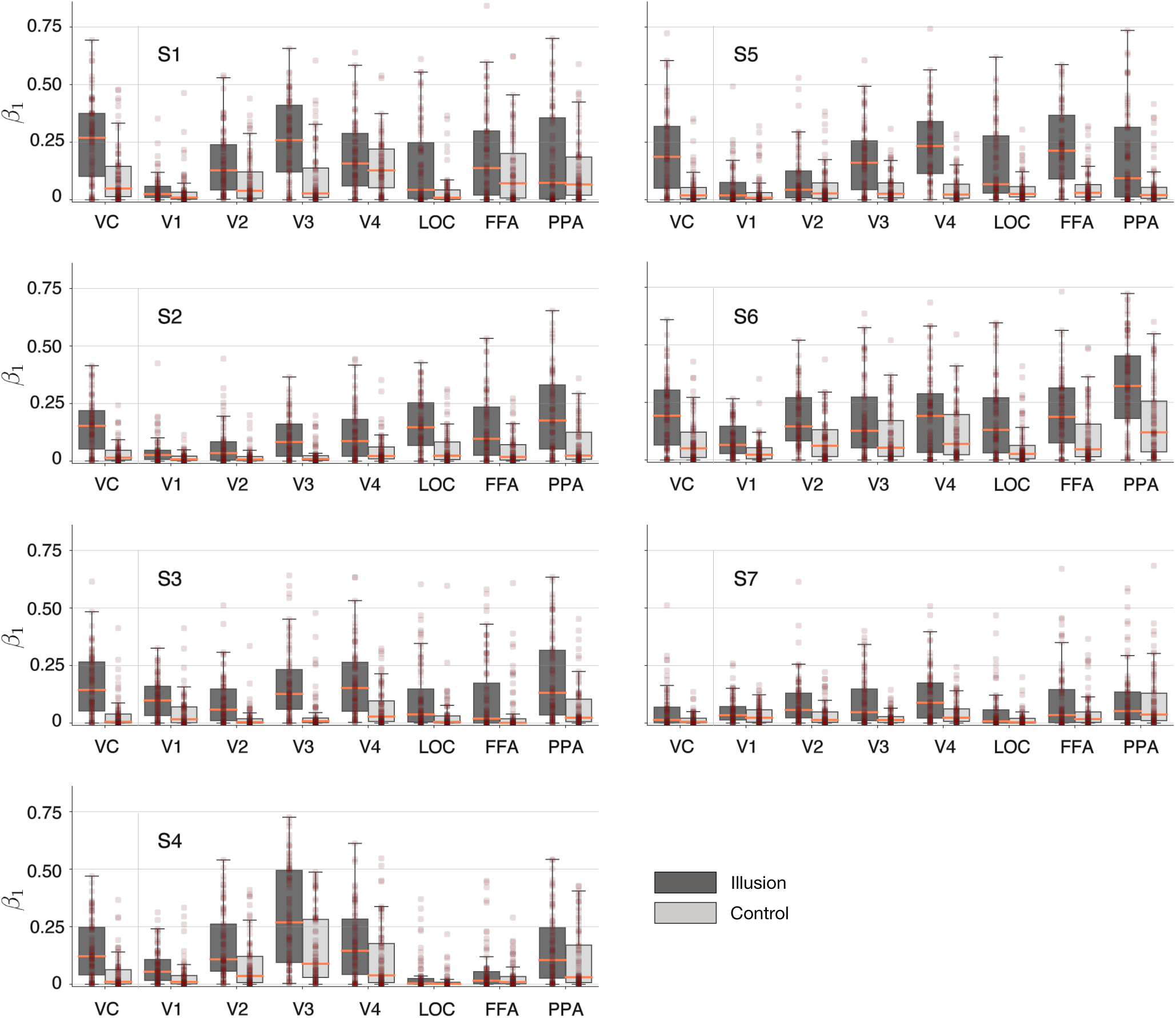
Comparison of the illusory surface coefficient values between illusion and control conditions of Ehrenstein for individual subjects. Results are based on single-trial reconstructions from VC and individual visual areas. Dots represent individual trials (the degree of opacity provides an indication of the density of dots). Coral lines show the median value and shaded areas of boxplots show the inter-quantile range of trials.

**Figure S21.**
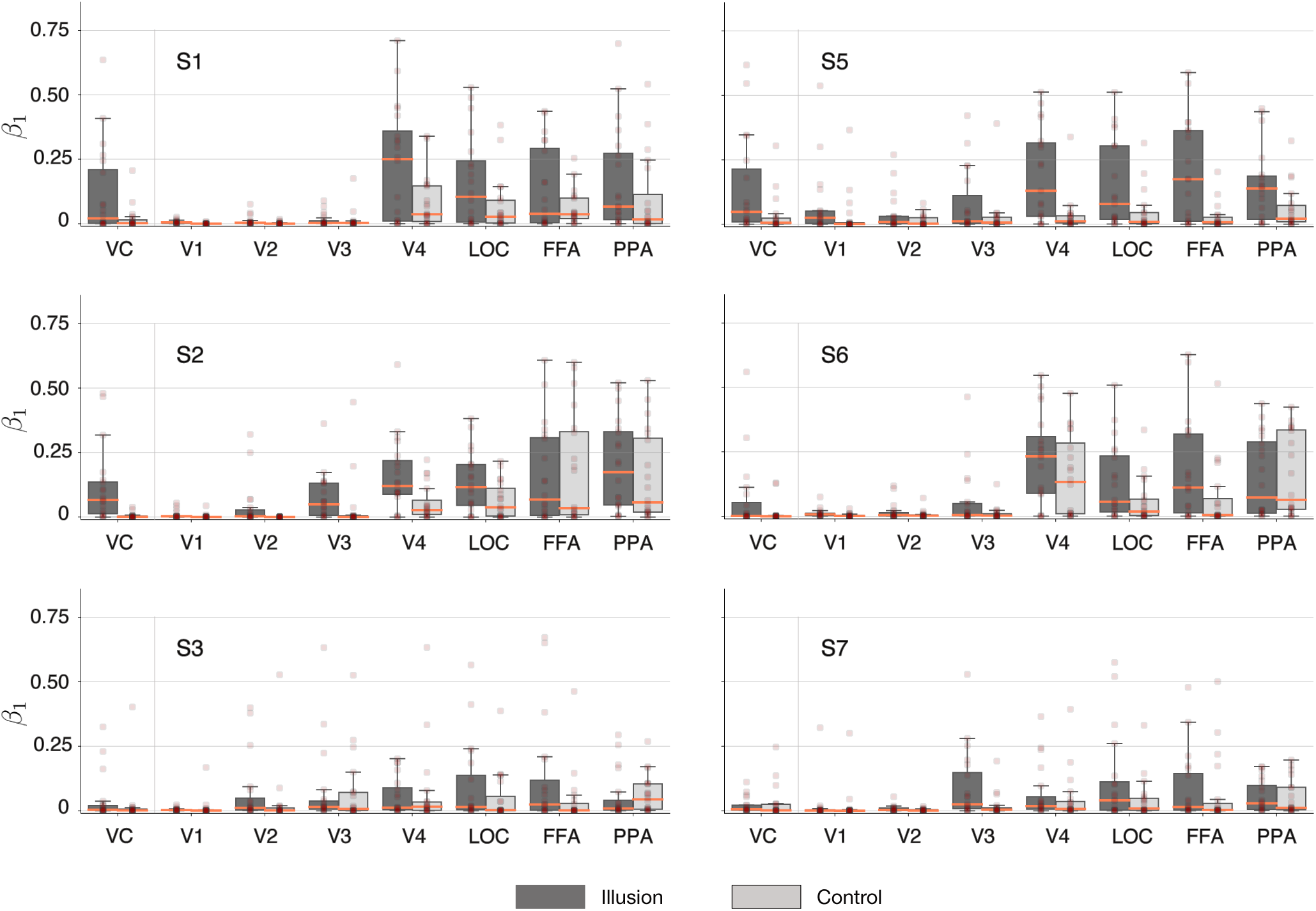
Comparison of the illusory surface coefficient values between illusion and control conditions of Varin for individual subjects. Results are based on single-trial reconstructions from VC and individual visual areas. Dots represent individual trials (the degree of opacity provides an indication of the density of dots). Coral lines show the median value and shaded areas of boxplots show the inter-quantile range of trials.

**Figure S22.**
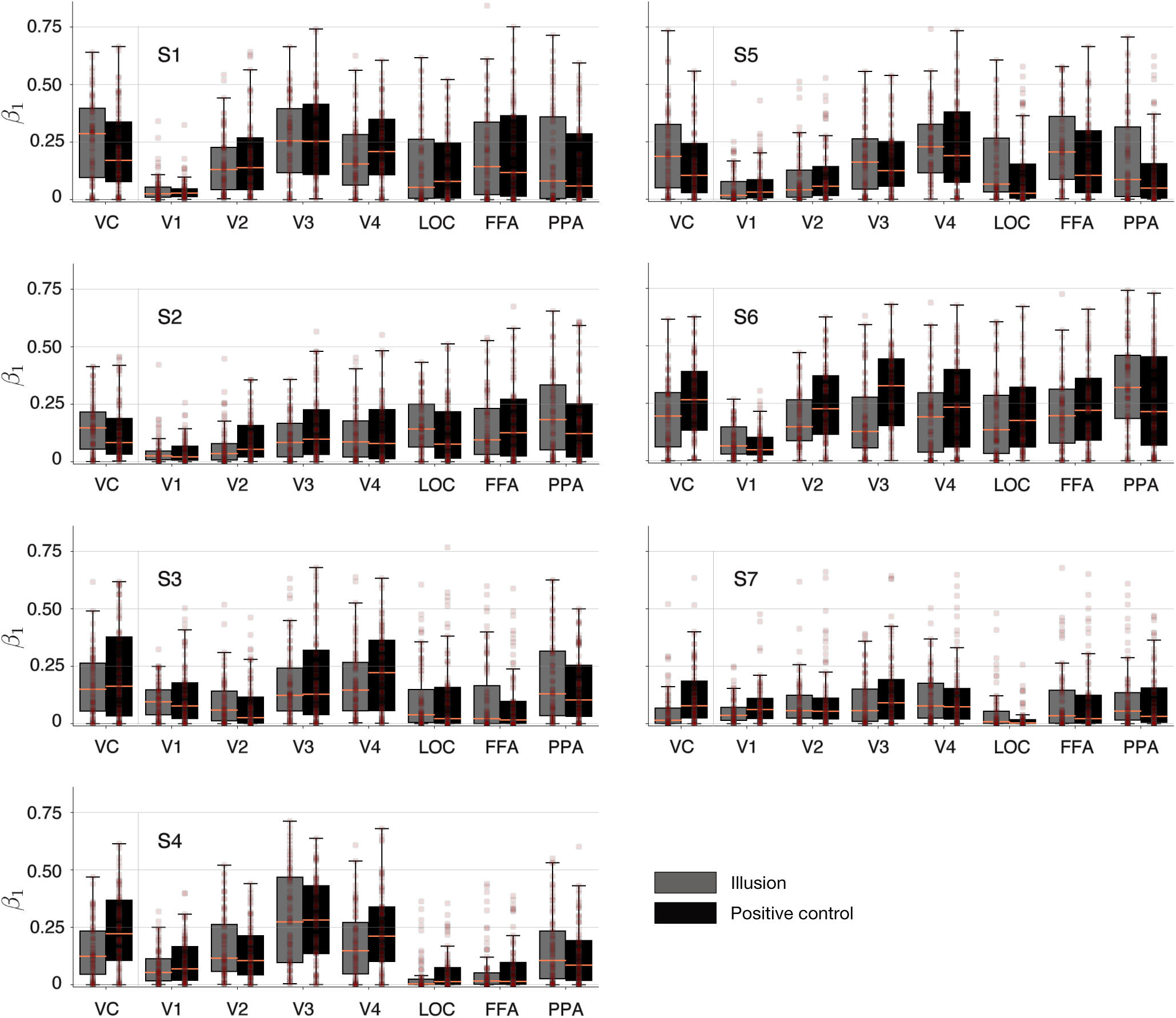
Comparison of the illusory surface coefficient values between illusion and positive control conditions of Ehrenstein for individual subjects. Results are based on single-trial reconstructions from VC and individual visual areas. Dots represent individual trials (the degree of opacity provides an indication of the density of dots). Coral lines show the median value and shaded areas of boxplots show the inter-quantile range of trials.

**Figure S23.**
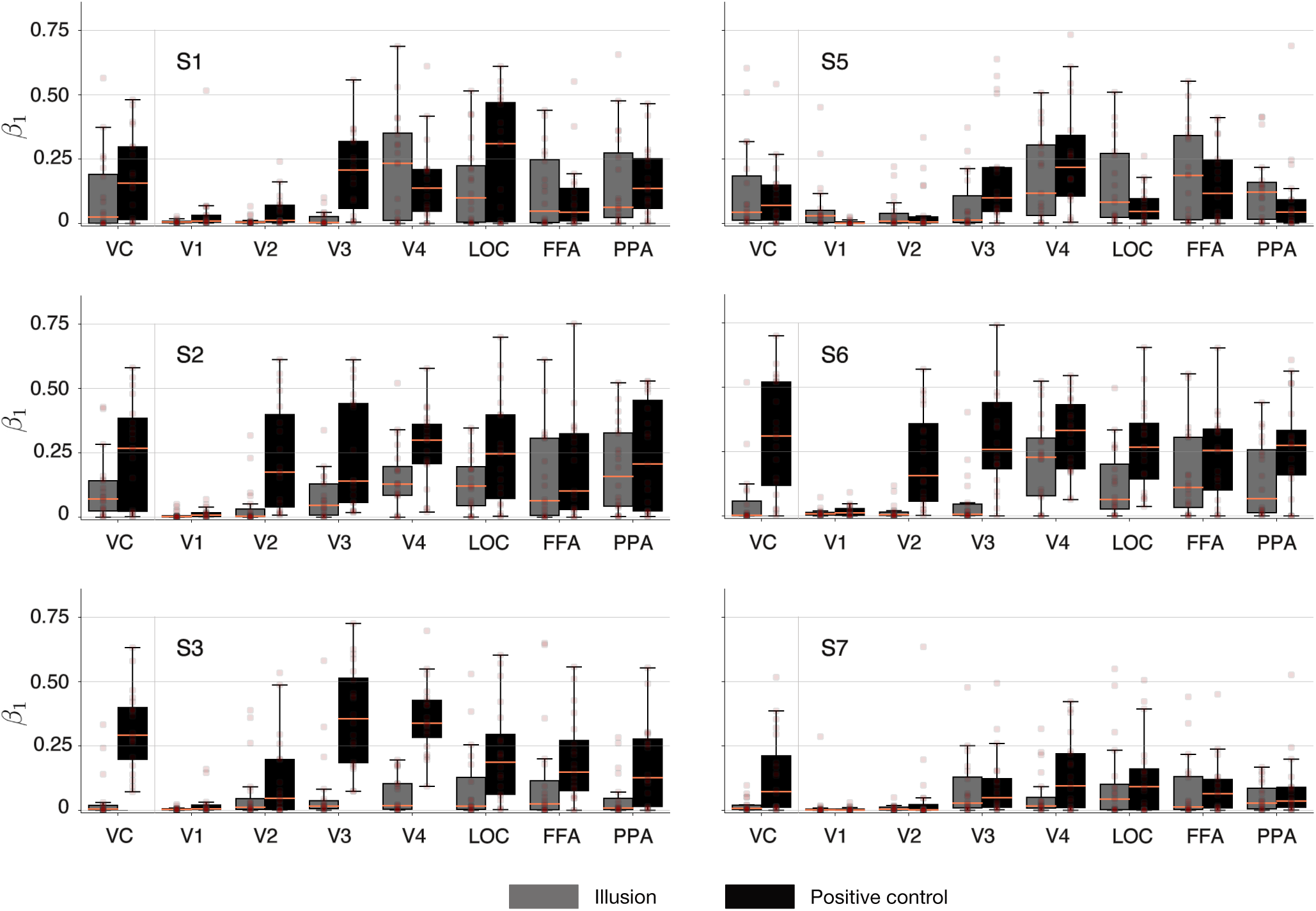
Comparison of the illusory surface coefficient values between illusion and positive control conditions of Varin for individual subjects. Results are based on single-trial reconstructions from VC and individual visual areas. Dots represent individual trials (the degree of opacity provides an indication of the density of dots). Coral lines show the median value and shaded areas of boxplots show the inter-quantile range of trials.

## Notes

### Competing Interest Statement

The authors have declared no competing interest.

### Summary of Updates

Figure 1 revised

## References

1. Bressan, P., Mingolla, E., Spillmann, L., and Watanabe, T. Neon Color Spreading: A Review. Perception, 26(11):1353–1366, Nov. 1997. doi: 10.1068/p261353.

2. Cornelissen, F. W., Wade, A. R., Vladusich, T., Dougherty, R. F., and Wandell, B. A. No Functional Magnetic Resonance Imaging Evidence for Brightness and Color Filling-In In Early Human Visual Cortex. Journal of Neuroscience, 26(14):3634–3641, Apr. 2006. doi: 10.1523/JNEUROSCI.4382-05.2006.

3. Cox, M. A., Schmid, M. C., Peters, A. J., Saunders, R. C., Leopold, D. A., and Maier, A. Receptive field focus of visual area V4 neurons determines responses to illusory surfaces. Proceedings of the National Academy of Sciences, 110(42):17095–17100, Oct. 2013. doi: 10.1073/pnas.1310806110.

4. Dhariwal, P. and Nichol, A. Diffusion Models Beat GANs on Image Synthesis. Machine Learning arXiv:2105.05233, June 2021.

5. Dosovitskiy, A. and Brox, T. Generating Images with Perceptual Similarity Metrics based on Deep Networks. In *Advances in Neural Information Processing Systems*, volume 29, 2016.

6. Ehrenstein, W. Über abwandlungen der l. hermannschen helligkeitserscheinung. Zeitschrift für Psychologie, 150:83–91, 1941.

7. Engel, S. A., Rumelhart, D. E., Wandell, B. A., Lee, A. T., Glover, G. H., Chichilnisky, E.-J., and Shadlen, M. N. fMRI of human visual cortex. Nature, 369(6481):525–525, June 1994. doi: 10.1038/369525a0.

8. Epstein, R. and Kanwisher, N. A cortical representation of the local visual environment. Nature, 392(6676):598–601, Apr. 1998. doi: 10.1038/33402.

9. Gerardin, P., Abbatecola, C., Devinck, F., Kennedy, H., Dojat, M., and Knoblauch, K. Neural circuits for long-range color filling-in. NeuroImage, 181:30–43, Nov. 2018. doi: 10.1016/j.neuroimage.2018.06.083.

10. Gomez-Villa, A., Martín, A., Vazquez-Corral, J., Bertalmío, M., and Malo, J. Color illusions also deceive CNNs for low-level vision tasks: Analysis and implications. Vision Research, 176:156–174, Nov. 2020. doi: 10.1016/j.visres.2020.07.010.

11. Grosof, D. H., Shapley, R. M., and Hawken, M. J. Macaque VI neurons can signal ‘illusory’ contours. Nature, 365(6446):550–552, Oct. 1993. doi: 10.1038/365550a0.

12. Heydt, R. v. d., Peterhans, E., and Baumgartner, G. Illusory contours and cortical neuron responses. Science, 224(4654):1260–1262, June 1984. doi: 10.1126/science.6539501.

13. Ho, M.-L. and Schwarzkopf, D. S. The human primary visual cortex (V1) encodes the perceived position of static but not moving objects. Communications Biology, 5(1):1–8, Mar. 2022. doi: 10.1038/s42003-022-03136-y.

14. Ho, J., Jain, A., and Abbeel, P. Denoising Diffusion Probabilistic Models. Machine Learning arXiv:2006.11239, Dec. 2020.

15. Hong, S. W. and Tong, F. Neural representation of form-contingent color filling-in in the early visual cortex. Journal of Vision, 17(13):10–10, Nov. 2017. doi: 10.1167/17.13.10.

16. Horikawa, T. and Kamitani, Y. Generic decoding of seen and imagined objects using hierarchical visual features. Nature Communications, 8:15037, May 2017. doi: 10.1038/ncomms15037.

17. Horikawa, T. and Kamitani, Y. Attention modulates neural representation to render reconstructions according to subjective appearance. Communications Biology, 5(1):1–12, Jan. 2022. doi: 10.1038/s42003-021-02975-5.

18. Ince, R. A. A., Kay, J. W., and Schyns, P. G. Within-participant statistics for cognitive science. Trends in Cognitive Sciences, June 2022. doi: 10.1016/j.tics.2022.05.008.

19. Jafari-Khouzani, K. and Soltanian-Zadeh, H. Radon transform orientation estimation for rotation invariant texture analysis. IEEE Transactions on Pattern Analysis and Machine Intelligence, 27(6):1004–1008, June 2005. doi: 10.1109/TPAMI.2005.126.

20. Kanwisher, N., McDermott, J., and Chun, M. M. The Fusiform Face Area: A Module in Human Extrastriate Cortex Specialized for Face Perception. The Journal of Neuroscience, 17 (11):4302–4311, June 1997. doi: 10.1523/JNEUROSCI.17-11-04302.1997.

21. Knebel, J.-F. and Murray, M. M. Towards a resolution of conflicting models of illusory contour processing in humans. NeuroImage, 59(3):2808–2817, Feb. 2012. doi: 10.1016/j.neuroimage.2011.09.031.

22. Kok, P. and de Lange, F. P. Shape Perception Simultaneously Upand Downregulates Neural Activity in the Primary Visual Cortex. Current Biology, 24(13):1531–1535, July 2014. doi: 10.1016/j.cub.2014.05.042.

23. Kok, P., Bains, L. J., van Mourik, T., Norris, D. G., and de Lange, F. P. Selective Activation of the Deep Layers of the Human Primary Visual Cortex by Top-Down Feedback. Current Biology, 26(3):371–376, Feb. 2016. doi: 10.1016/j.cub.2015.12.038.

24. Kourtzi, Z. and Kanwisher, N. Cortical Regions Involved in Perceiving Object Shape. The Journal of Neuroscience, 20(9):3310–3318, May 2000. doi: 10.1523/JNEUROSCI.20-09-03310.2000.

25. Krizhevsky, A., Sutskever, I., and Hinton, G. E. ImageNet Classification with Deep Convolutional Neural Networks. In *Advances in Neural Information Processing Systems*, volume 25, 2012.

26. Lee, T. S. and Nguyen, M. Dynamics of subjective contour formation in the early visual cortex. Proceedings of the National Academy of Sciences, 98(4):1907–1911, Feb. 2001. doi: 10.1073/pnas.98.4.1907.

27. Lin, T.-Y., Maire, M., Belongie, S., Bourdev, L., Girshick, R., Hays, J., Perona, P., Ramanan, D., Zitnick, C. L., and Dollár, P. Microsoft COCO: Common Objects in Context. May 2014. doi: 10.48550/arXiv.1405.0312.

28. Lotter, W., Kreiman, G., and Cox, D. A neural network trained for prediction mimics diverse features of biological neurons and perception. Nature Machine Intelligence, 2(4):210– 219, Apr. 2020. doi: 10.1038/s42256-020-0170-9.

29. Metelli, F. Stimulation and perception of transparency. Psychological Research, 47(4):185–202, Dec. 1985. doi: 10.1007/BF00309446.

30. Miyawaki, Y., Uchida, H., Yamashita, O., Sato, M.-a., Morito, Y., Tanabe, H. C., Sadato, N., and Kamitani, Y. Visual Image Reconstruction from Human Brain Activity using a Combination of Multiscale Local Image Decoders. Neuron, 60(5):915–929, Dec. 2008. doi: 10.1016/j.neuron.2008.11.004.

31. Nakayama, K., Shimojo, S., and Ramachandran, V. S. Transparency: Relation to Depth, Subjective Contours, Luminance, and Neon Color Spreading. Perception, 19(4):497– 513, Aug. 1990. doi: 10.1068/p190497.

32. Pak, A., Ryu, E., Li, C., and Chubykin, A. A. Top-Down Feedback Controls the Cortical Representation of Illusory Contours in Mouse Primary Visual Cortex. Journal of Neuroscience, Dec. 2019. doi: 10.1523/JNEUROSCI.1998-19.2019.

33. Pan, Y., Chen, M., Yin, J., An, X., Zhang, X., Lu, Y., Gong, H., Li, W., and Wang, W. Equivalent Representation of Real and Illusory Contours in Macaque V4. Journal of Neuroscience, 32(20):6760–6770, May 2012. doi: 10.1523/JNEUROSCI.6140-11.2012.

34. Peterhans, E. and Heydt, R. v. d. Mechanisms of contour perception in monkey visual cortex. II. Contours bridging gaps. Journal of Neuroscience, 9(5):1749–1763, May 1989. doi: 10.1523/JNEUROSCI.09-05-01749.1989.

35. Ramsden, B. M., Hung, C. P., and Roe, A. W. Real and Illusory Contour Processing in Area V1 of the Primate: a Cortical Balancing Act. Cerebral Cortex, 11(7):648–665, July 2001. doi: 10.1093/cercor/11.7.648.

36. Redies, C. and Spillmann, L. The Neon Color Effect in the Ehrenstein Illusion. Perception, 10(6):667–681, Dec. 1981. doi: 10.1068/p100667.

37. Roe, A. W., Lu, H. D., and Hung, C. P. Cortical processing of a brightness illusion. Proceedings of the National Academy of Sciences, 102(10):3869–3874, Mar. 2005. doi: 10.1073/pnas.0500097102.

38. Russakovsky, O., Deng, J., Su, H., Krause, J., Satheesh, S., Ma, S., Huang, Z., Karpathy, A., Khosla, A., Bernstein, M., Berg, A. C., and Fei-Fei, L. ImageNet Large Scale Visual Recognition Challenge. International Journal of Computer Vision, 115(3):211–252, Dec. 2015. doi: 10.1007/s11263-015-0816-y.

39. Saeedi, A., Wang, K., Nikpourian, G., Bartels, A., Totah, N. K., Logothetis, N. K., and Watan-abe, M. Mouse primary visual cortex neurons respond to the illusory “darker than black” in neon color spreading. bioRxiv, 2022. doi: 10.1101/2022.07.24.501311.

40. Sáry, G., Chadaide, Z., Tompa, T., Köteles, K., Kovács, G., and Benedek, G. Illusory shape representation in the monkey inferior temporal cortex. European Journal of Neuroscience, 25(8):2558–2564, 2007. doi: 10.1111/j.1460-9568.2007.05494.x.

41. Sasaki, Y. and Watanabe, T. The primary visual cortex fills in color. Proceedings of the Na-tional Academy of Sciences, 101(52):18251–18256, Dec. 2004. doi: 10.1073/pnas.0406293102.

42. Schumann, F. Beiträge zur analyse der gesichtswahrnehmungen. erste abhandlung. einige beobachtungen über die zusammenfassung von gesichtseindrücken zu einheiten. Zeitschrift für Psychologie und Physiologie der Sinnesorgane, 23:1–32, 1900.

43. Seghier, M. L. and Vuilleumier, P. Functional neuroimaging findings on the human perception of illusory contours. Neuroscience & Biobehavioral Reviews, 30(5):595–612, Jan. 2006. doi: 10.1016/j.neubiorev.2005.11.002.

44. Sereno, M., Dale, A., Reppas, J., Kwong, K., Belliveau, J., Brady, T., Rosen, B., and Tootell, R. Borders of multiple visual areas in humans revealed by functional magnetic resonance imaging. Science, 268(5212):889–893, May 1995. doi: 10.1126/science.7754376.

45. Sharan, L., Rosenholtz, R., and Adelson, E. H. Accuracy and speed of material categorization in real-world images. Journal of Vision, 14(9):12, Aug. 2014. doi: 10.1167/14.9.12.

46. Shen, G., Horikawa, T., Majima, K., and Kamitani, Y. Deep image reconstruction from human brain activity. PLOS Computational Biology, 15(1):1006633, Jan. 2019a. doi: 10.1371/journal.pcbi.1006633.

47. Shen, G., Dwivedi, K., Majima, K., Horikawa, T., and Kamitani, Y. End-to-End Deep Image Reconstruction From Human Brain Activity. Frontiers in Computational Neuroscience, 13:21, Apr. 2019b. doi: 10.3389/fncom.2019.00021.

48. Soriano, M., Spillmann, L., and Bach, M. The abutting grating illusion. Vision Research, 36 (1):109–116, Jan. 1996. doi: 10.1016/0042-6989(95)00107-B.

49. Sun, E. D. and Dekel, R. ImageNet-trained deep neural networks exhibit illusion-like response to the Scintillating grid. Journal of Vision, 21(11):15, Oct. 2021. doi: 10.1167/jov.21.11.15.

50. van Tuijl, H. F. J. M. A new visual illusion: Neonlike color spreading and complementary color induction between subjective contours. Acta Psychologica, 39(6):441–445, Dec. 1975. doi: 10.1016/0001-6918(75)90042-6.

51. VanRullen, R. and Reddy, L. Reconstructing faces from fMRI patterns using deep genera-tive neural networks. Communications Biology, 2(1):1–10, May 2019. doi: 10.1038/s42003-019-0438-y.

52. Varin, D. Fenomini di contrasto e diffusione cromatica nell’organizzazione spaziale del campo percettivo. Rivista di Psicologia, 65:101–128, 1971.

53. Watanabe, E., Kitaoka, A., Sakamoto, K., Yasugi, M., and Tanaka, K. Illusory Motion Reproduced by Deep Neural Networks Trained for Prediction. Frontiers in Psychology, 9:345. doi: 10.3389/fpsyg.2018.00345, 2018.

54. Yang, L., Zhang, Z., Song, Y., Hong, S., Xu, R., Zhao, Y., Zhang, W., Cui, B., and Yang, M.-H. Diffusion Models: A Comprehensive Survey of Methods and Applications. Machine Learning arXiv:2209.00796, Mar. 2023.

